# Orbitofrontal control of visual cortex gain promotes visual associative learning

**DOI:** 10.1101/794958

**Authors:** Dechen Liu, Juan Deng, Zhewei Zhang, Zhi-Yu Zhang, Yan-Gang Sun, Tianming Yang, Haishan Yao

## Abstract

The orbitofrontal cortex (OFC) encodes expected outcomes and plays a critical role in flexible, outcome-guided behavior. The OFC projects to primary visual cortex (V1), yet the function of this top-down projection is unclear. We find that optogenetic activation of OFC projection to V1 reduces the amplitude of V1 visual responses via the recruitment of local somatostatin-expressing (SST) interneurons. Using mice performing a Go/No-Go visual task, we show that the OFC projection to V1 mediates the outcome-expectancy modulation of V1 responses to the reward-irrelevant No-Go stimulus. Furthermore, V1-projecting OFC neurons reduce firing during expectation of reward. In addition, chronic optogenetic inactivation of OFC projection to V1 impairs, whereas chronic activation of SST interneurons in V1 improves the learning of Go/No-Go visual task, without affecting the immediate performance. Thus, OFC top-down projection to V1 is crucial to drive visual associative learning by modulating the response gain of V1 neurons to non-relevant stimulus.

---

The OFC is involved in encoding specific stimulus-reward associations that guide adaptive behavior^1-3^. Studies in rodents and monkeys have demonstrated that the identity and expected values of specific outcomes are represented by activities in the OFC^4-12^. Lesions or inactivation of the OFC impair behavior guided by outcome expectancy and learning driven by the discrepancy between expected and actual outcomes^13-26^, and degrade the acquisition of Pavlovian trace conditioning task^27^. Direct output from the OFC to other brain regions, including the basolateral amygdala (BLA), ventral tegmental area (VTA) and striatum, is important for learning and reward-related behavior^2,3^. In OFC-lesioned rats, BLA neurons are impaired in the encoding of cue-outcome association and the developing of outcome-expectant activity^28^. The outcome-expectancy signals in the OFC are necessary for VTA dopamine neurons to calculate reward prediction errors^29^, which are important teaching signals for reinforcement learning^30^. Inactivation of the OFC or disconnection of the OFC from VTA prevents extinction learning in the Pavlovian over-expectation task^23,31^. The VTA-projecting OFC neurons encode long-term memory of cue-reward association, and optogenetic inhibition of these neurons impairs extinction learning and memory^27^. The OFC also connects with sensory cortices^32-34^, including V1^32^. It is unknown how the responses of sensory cortex-projecting OFC neurons are modulated by outcome expectancy, and whether the top-down signals from the OFC to sensory cortices influence learning behavior.

Frontal top-down projections to sensory cortices are known to modulate sensory processing^35-37^, promote accurate perception^38^, and convey predictive signals^39,40^. Associative learning enhances signals related to stimulus expectation or reward expectation in V1, which may be mediated by top-down projections^41^. Learning also enhances the effect of top-down inputs in modulating V1 responses^42^. However, the causal role of top-down projections to sensory cortices in stimulus-reward associative learning remains unclear. In this study, we find that activation of OFC top-down projection results in suppression of V1 visual response by activating SST interneurons. In mice performing a Go/No-Go visual task, V1 responses to the reward-irrelevant No-Go stimulus are lower when the mice’ outcome expectation is correct than when it is incorrect, and such response modulation is reduced by optogenetic inactivation of OFC projection to V1. Optogenetic tagging of V1-projecting OFC neurons reveals that their responses to the No-Go stimulus are reduced in trials in which mice performed incorrectly. We further show that chronic optogenetic inactivation of OFC projection to V1 slows the learning of Go/No-Go visual behavior. Thus, the OFC projection to V1 plays a key role in filtering out non-relevant visual information to facilitate associative learning.

## Results

### Effect of activating OFC top-down projection on V1 responses

We injected Cholera toxin subunit B (CTB) in V1 and found that the retrograde labeled neurons were in the ventrolateral OFC (vlOFC) (Fig. 1a), consistent with the finding in a previous study^32^. By injecting rAAV2-retro-hSyn-Cre in V1 and AAV-DIO-EYFP in the OFC, we found that OFC axons terminated in both superficial and deep layers of V1 (Fig. 1b). To examine how OFC top-down projection influences V1 neuronal responses, we expressed excitatory opsin Channelrhodopsin-2 (ChR2) or ChrimsonR in the OFC, and measured V1 responses with and without laser stimulation of OFC axons in mice passively viewing drifting gratings (Fig. 1c). Activating OFC axons in V1 significantly reduced the firing rates of V1 neurons in both anesthetized and awake mice (anesthetized mice: *P* = 5.2×10^-6^, *n* = 102 neurons; awake mice: *P* = 6.53×10^-5^, *n* = 62 neurons; Wilcoxon signed rank test, Fig. 1d). When we computed a rate change index as (R_laser_on_ - R_laser_off_)/(R_laser_on_ + R_laser_off_), where R_laser_on_ and R_laser_off_ represented responses averaged over all orientations for laser-on and laser-off trials, respectively, we found that the index was negative for the majority of V1 neurons in both anesthetized and awake mice (Fig. 1e). For control mice injected with AAV-mCherry (or AAV-EGFP) in the OFC, the laser-induced response reduction was significantly smaller than that for mice injected with AAV-ChR2 (or AAV-ChrimsonR) (Supplementary Fig. 1a). After blocking antidromic spikings of OFC neurons with tetrodotoxin in the OFC, the response reduction in V1 neurons induced by activating OFC axons was still significant (*P* = 2.33×10^-12^, *n* = 118 neurons from awake mice, Wilcoxon signed rank test, Fig. 1f−h), indicating that the laser-induced response reduction was mediated directly by OFC projection to V1, rather than through antidromic activation of indirect pathways. Although the response amplitude was clearly reduced, the orientation selectivity of V1 neurons was not affected by activation of OFC axons in V1 (anesthetized mice: *P* = 0.36, *n* = 102 neurons; awake mice: *P* = 0.99, *n* = 62 neurons; Wilcoxon signed rank test, Fig. 1i).

**Fig. 1.**
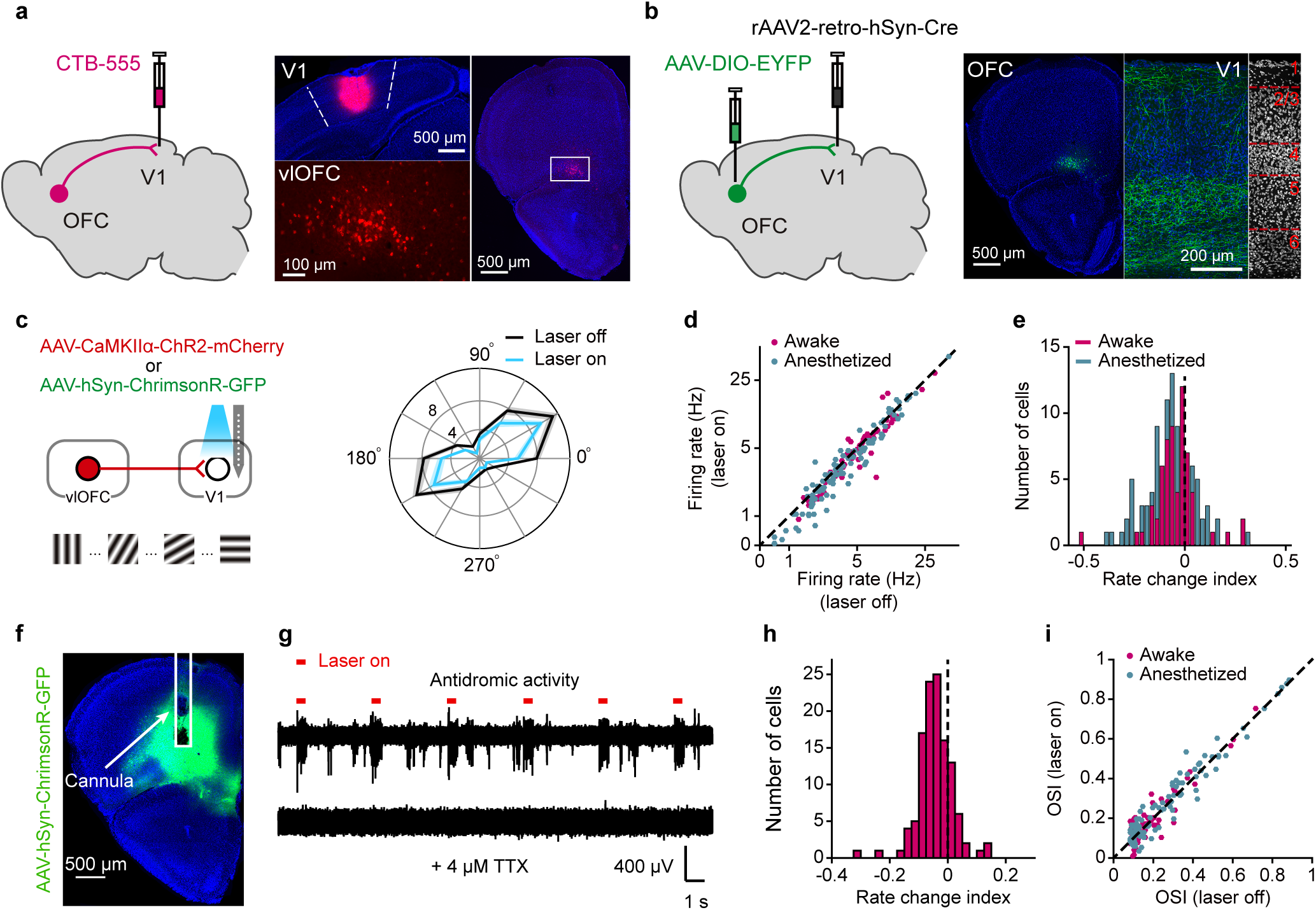
Activating OFC projection to V1 reduces response amplitude of V1 neurons. **a**, Left, schematic of CTB injection in V1. Right, Representative fluorescence images of CTB injection in V1 and retrograde labeled neurons in the ventrolateral OFC. **b**, Left, strategy of virus injection to visualize the OFC axons in V1. Right, representative fluorescence images of V1-projecting OFC neurons and their terminals in V1. **c**, Left, schematic of measuring V1 visual responses with and without activating OFC axons in V1. Right, tuning curves of a V1 neuron with (blue) and without (black) laser stimulation of OFC axons in V1. **d**, Mean firing rate (firing rate averaged over all orientations) of V1 neurons with laser on vs. laser off. Anesthetized mice (blue): *P* = 5.2×10^-6^, *n* = 102 neurons; awake mice (magenta): *P* = 6.53×10^-5^, *n* = 62 neurons. **e**, Distribution of rate change indexes for V1 neurons in anesthetized (*P* = 2.6×10^-8^) and awake mice (*P* = 3.81×10^-5^). The rate change index was computed as (R_laser_on_ - R_laser_off_)/(R_laser_on_ + R_laser_off_), in which R_laser_on_ and R_laser_off_ represented responses averaged over all orientations for laser-on and laser-off trials, respectively. **f**, TTX was infused to the OFC, in which AAV-hSyn-ChrimsonR-GFP had been injected, to block the antidromic spikes induced by laser stimulation in V1. White rectangle shows the placement of the cannula. **g**, TTX infusion into the OFC abolished multi-unit activity in the OFC evoked by laser stimulation in V1. Upper trace, without TTX; Lower trace, with TTX infusion into the OFC. **h**, Distribution of rate change indexes of V1 neurons recorded with TTX infusion into the OFC. *P* = 2.33×10^-12^*, n* = 118 neurons from awake mice. **i**, Orientation selectivity index (OSI) with laser on vs. laser off. Anesthetized mice (blue): *P* = 0.36, *n* = 102; awake mice (magenta): *P* = 0.99, *n* = 62. For **d**, **e**, **h** and **i**, AAV-CaMKIIα-hChR2 (H134R)-mCherry and AAV-hSyn-ChrimsonR-GFP were injected in the OFC for anesthetized and awake mice, respectively. Wilcoxon two-sided signed rank test. For **d**, **e**, **h** and **i**, source data are provided as a Source Data file. Shadings, mean ± s.e.m.

To dissect the circuit mechanism underlying V1 response modulation by the OFC top-down projection, we infected the OFC neurons with AAV-ChR2 and performed whole-cell recordings from V1 neurons in acute slices of V1 containing ChR2-expressing OFC axons. Photostimulation of OFC axons evoked both excitatory and inhibitory postsynaptic currents (EPSCs and IPSCs) in the recorded neurons (Fig. 2a). The onset latencies of the EPSCs were shorter than those of the IPSCs (Fig. 2b), which were blocked by γ-aminobutyric acid type A (GABA_A_) receptor antagonist picrotoxin (Fig. 2c). We next tested whether IPSCs evoked by activation of OFC axons were feedforward inhibition. We bath applied an α-amino-3-hydroxy-5-methyl-4-isoxazole propionic acid (AMPA) receptor antagonist NBQX, and found that the amplitudes of both EPSCs and IPSCs were reduced (Fig. 2d), indicating that the IPSCs were due to feedforward inhibition generated by local inhibitory neurons in V1. By injecting CTB in V1 of GAD67-GFP mice, we found that the retrograde labeling of OFC neurons did not overlap with the GFP-positive neurons in the OFC (Fig. 2e), confirming that the V1-projecting OFC neurons were not GABAergic.

**Fig. 2.**
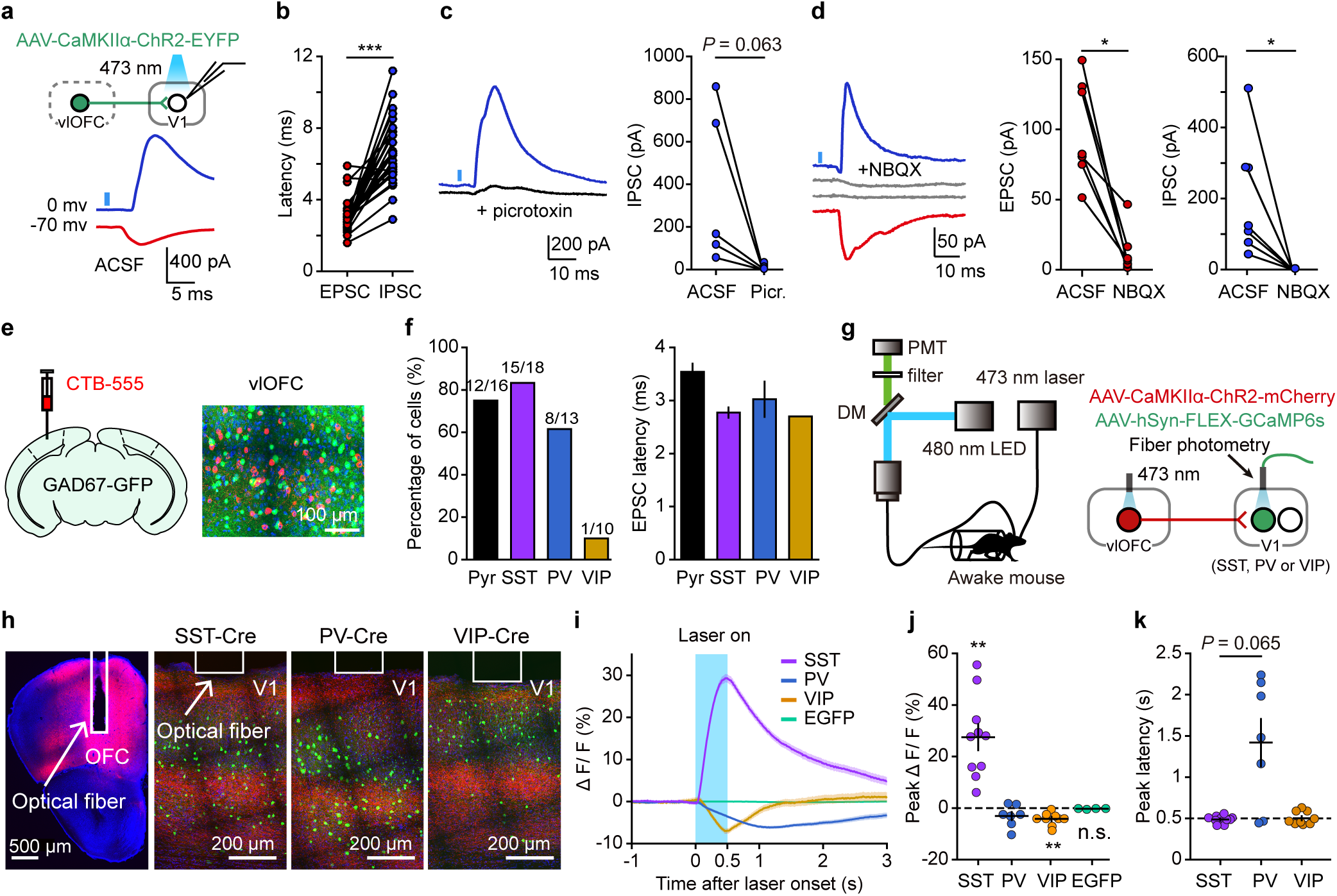
Optogenetic activation of OFC axons activates inhibitory interneurons in V1. **a**, Photostimulation of OFC projection to V1 evoked both EPSCs (red) and IPSCs (blue) in a V1 neuron. **b**, The onset latencies of laser-evoked EPSCs were significantly shorter than those of IPSCs (\*\*\**P* = 2.34×10^-6^, *n* = 30, in which *n* = 7, 12, 1, 1 and 9 for pyramidal, SST, PV, VIP and unidentified neurons, respectively). **c**, Left, application of a GABA_A_ receptor antagonist (picrotoxin, 30 μM) blocked IPSCs in a V1 neuron. Black, IPSCs after the application of picrotoxin. Right, comparison of IPSCs amplitudes before and after picrotoxin application (*P* = 0.063, *n* = 5). **d**, Left, application of an AMPA receptor antagonist (NBQX, 10 μM) blocked both EPSCs and IPSCs in a V1 neuron. Gray, EPSCs and IPSCs after the application of NBQX. Middle and right, comparison of EPSCs (IPSCs) amplitudes before and after NBQX application (*P* = 0.016 and *n* = 7 for both EPSCs and IPSCs). **e**, CTB was injected in V1 of a GAD67-GFP mouse. Retrograde labeling of OFC neurons did not overlap with GFP-positive neurons in the OFC. The experiments were repeated in 3 mice. **f**, Left, fraction of pyramidal (Pyr) and inhibitory interneurons in which EPSCs could be evoked by laser stimulation of OFC axons in V1. Right, latency of EPSC for each type of neuron. **g**, Schematic of the strategy to record activities of specific interneuron type in V1 evoked by OFC activation *in vivo*. **h**, Fluorescence image of OFC in which AAV-CaMKIIα-ChR2-mCherry was injected, and those of V1 injected with AAV-hSyn-FLEX-GCaMP6s from a SST-Cre, PV-Cre and VIP-Cre mouse, respectively. White rectangles show the placement of the optic fiber. **i**, Representative ΔF/F signals from a SST-Cre, PV-Cre, VIP-Cre mouse and a C57BL/6 mouse (AAV-hSyn-EGFP injected in V1), respectively. Blue area, period of OFC stimulation. **j**, Amplitude of peak ΔF/F. SST: \*\**P* = 0.002, *n* = 10 mice; PV: *P* = 0.078, *n* = 7 mice; VIP: \*\**P* = 0.004, *n* = 9 mice; EGFP: *P* = 0.13, *n* = 4 mice. **k**, Latency of peak ΔF/F. SST vs PV: *P* = 0.065 (Wilcoxon two-sided rank sum test). Wilcoxon two-sided signed rank test for **b**, **c**, **d** and **j**. For **b**−**d**, **f** and **i**−**k**, source data are provided as a Source Data file. Shadings and error bars, mean ± s.e.m.

We next examined which subtype of inhibitory neurons mediates the effect of activating OFC axons on V1 responses. By using rabies virus (RV)-mediated monosynaptic retrograde tracing^43^, we found that V1 interneurons expressing parvalbumin (PV), SST and vasoactive intestinal peptide (VIP) all received direct innervation from neurons in the vlOFC (Supplementary Fig. 2). Consistently, slice recording showed that optogenetic stimulation of OFC axons evoked EPSCs in three different subtypes of interneurons as well as pyramidal neurons in V1 (Fig. 2f). As the latencies of EPSCs were short (Fig. 2f), the EPSCs were likely evoked by direct excitatory drive from the OFC.

We further used fiber photometry to measure the activities of three subtypes of V1 interneuron *in vivo* in response to optogenetic activation of OFC neurons (Fig. 2g). For these mice, we expressed ChR2 in the OFC and calcium indicator GCaMP6s in PV, SST, or VIP interneurons in V1, respectively (Fig. 2h). During the experiment, the mice were awake but were not viewing visual stimulus. We found that optogenetic activation of OFC neurons caused increase in calcium signals in SST interneurons, but reduction in calcium signals in PV and VIP interneurons in V1 (Fig. 2i, j). Since the latency of laser-evoked peak responses for PV interneurons was longer than that for SST interneurons (Fig. 2k), the reduced PV neuronal responses were likely to be attributed to inhibition caused by SST interneuron activation^44^. Early activated population of SST interneurons could also inhibit VIP interneurons^44^, causing their activity reduction. Therefore, OFC stimulation *in vivo* preferentially activated SST interneurons in V1, providing a circuit mechanism for top-down modulation of V1 responses by the OFC.

### OFC projection modulates V1 responses to irrelevant stimulus

To examine whether inactivating the OFC projection to V1 influences V1 neuronal responses, we expressed inhibitory opsin Jaws in the OFC. By recording V1 neurons from awake mice passively viewing drifting gratings, we found that inactivating the OFC projection to V1 did not cause significant change in V1 responses as compared to the control mice (*P* = 0.17, Wilcoxon rank sum test, Supplementary Fig. 1b, c).

We next wondered whether the OFC projection to V1 may function during task engagement to suppress V1 responses to non-relevant visual stimulus. To test this hypothesis, we trained head-fixed mice to perform a Go/No-Go visual task (Fig. 3a, b), in which a vertical grating (the “Go” stimulus) and a horizontal grating (the “No-Go” stimulus) were associated with water reward and no reward, respectively. In each trial, the duration of stimulus presentation included a waiting period, during which licking had no consequence, and an answer period (Fig. 3a). For a Go trial, licking within the answer period was rewarded with water (hit). For a No-Go trial, licking (false alarm, FA) within the answer period was neither rewarded nor punished, and withholding licking within the answer period represented correct rejection (CR). During the inter-trial interval (ITI), the screen was blank and licking was punished with a longer ITI. The training of the Go/No-Go task was preceded by 2 sessions of conditioning, in which the mouse learned to lick within the answer period after the presentation of a Go stimulus. We found that the latency of the first lick after stimulus onset increased with training, and the ITI decreased over sessions (Fig. 3b, Supplementary Fig. 3), indicating that the mice gradually understood the task structure. Over 11 training sessions of the Go/No-Go task, the hit rate remained high throughout all sessions, whereas the CR rate and discriminability (d’) increased with training (Fig. 3c, d). Thus, the learning of the task depended on the improvement of CR for the reward-irrelevant No-Go stimulus.

**Fig. 3.**
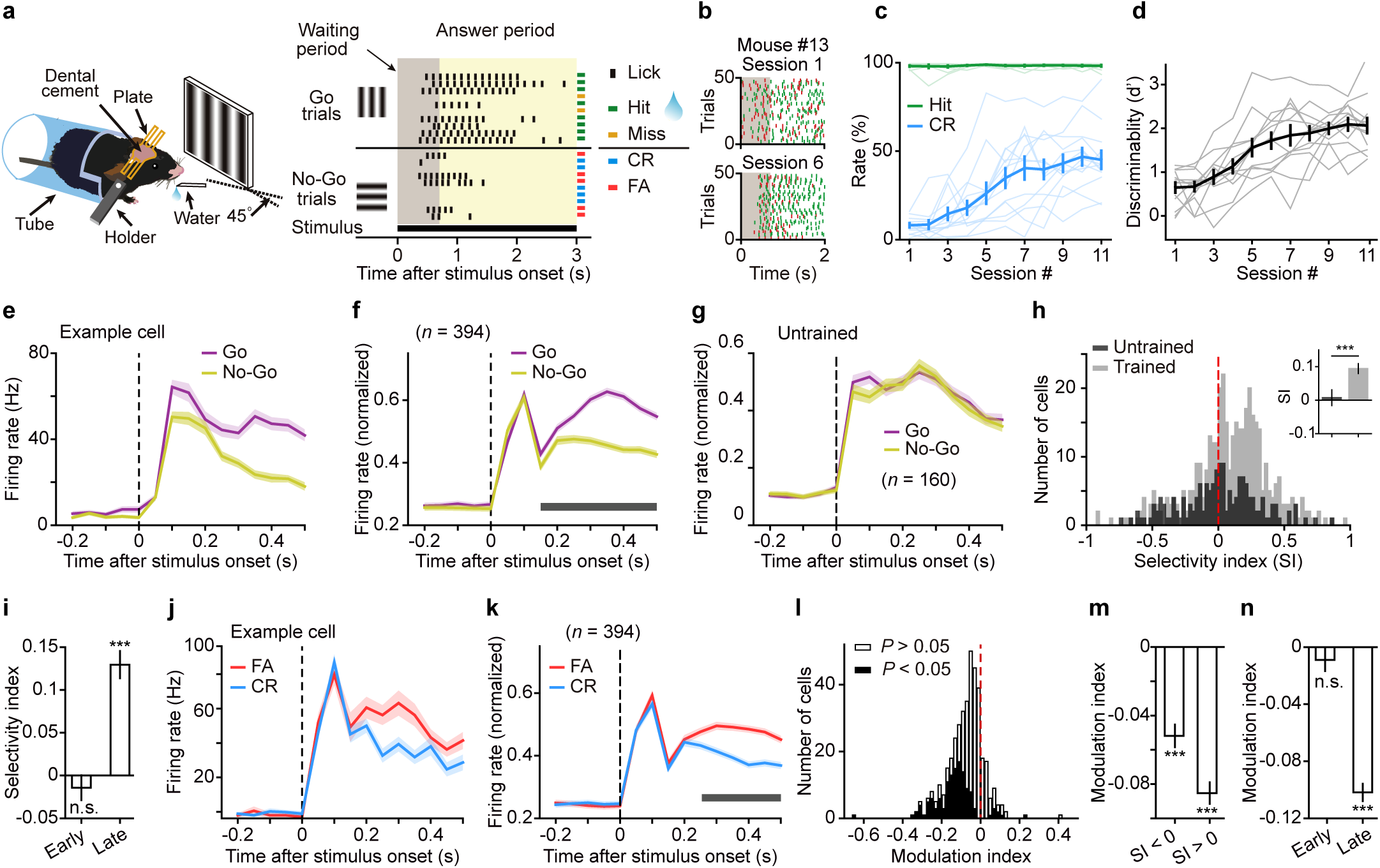
Responses of V1 neurons in Go/No-Go visual task. **a**, Left, behavioral setup. Right, schematic of task structure. In each trial, the duration of stimulus presentation included a waiting period (gray) and an answer period (yellow). CR, correct rejection. FA, false alarm. **b**, Lick rasters from a subset of trials in two behavioural sessions for a mouse. **c**, Hit (green) and CR rates (blue) over sessions (*n* = 13 mice). Each thin line represents a mouse. For hit rate, *F*_(2.72, 32.62)_ = 0.43, *P* = 0.71; for CR rate, *F*_(3.36, 40.28)_ = 23.57, *P* = 2.1×10^-9^. **d**, Discriminability over sessions. Each thin line represents a mouse. *F*_(4.49, 53.86)_ = 33.34, *P* = 1.1×10^-14^. One-way repeated measures ANOVA with the Greenhouse-Geisser correction. **e**, Responses of an example V1 neuron to the Go and No-Go stimuli. **f**, The firing rates of each V1 neuron were normalized by the maximum of the peak values in the Go and No-Go trials, and were averaged across neurons. Horizontal bar indicates time points in which the responses between Go and No-Go trials were significantly different (*P* < 0.05, two-way repeated measures ANOVA *F*_(1, 393)_ = 55.7, *P* = 5.5×10^-13^ followed by Sidak’s multiple comparisons test). **g**, Responses of V1 neurons from untrained mice passively viewing the stimulus (*P* = 0.56, two-way repeated measures ANOVA *F*_(1, 159)_ = 0.33). **h**, Distribution of selectivity indexes for V1 neurons. Light gray, trained mice, *P* = 2.2×10^-12^, *n* = 394 neurons. Dark gray, untrained mice, *P* = 0.62, *n* = 160 neurons. Wilcoxon two-sided signed rank test. Inset, comparison of SIs between V1 neurons in untrained and trained mice.\*\*\**P* = 4.6×10^-4^, Wilcoxon two-sided rank sum test. **i**, Selectivity index for early and late response components for V1 neurons from trained mice. Early component, *P* = 0.51; late component, \*\*\**P* = 2.1×10^-16^, *n* = 394, Wilcoxon two-sided signed rank test. **j**, Responses of an example V1 neuron to the No-Go stimulus in FA and CR trials. **k**, The firing rates of each V1 neuron to the No-Go stimulus were normalized by the maximum of the peak values in FA and CR trials, and were averaged across neurons. Horizontal bar indicates time points in which the responses between FA and CR conditions were significantly different (*P* < 0.05, two-way repeated measures ANOVA *F*_(1, 393)_ = 222.22, *P* < 1×10^-15^ followed by Sidak’s multiple comparisons test). **l**, Distribution of modulation indexes (MIs) for V1 neurons (*P* = 1.46×10^-42^, *n* = 394 neurons, Wilcoxon two-sided signed rank test). Open bar, MI was not significant. Black bar, MI was significant. **m**, The MIs for V1 neurons preferring the Go stimulus (SI > 0, *n* = 270) and preferring the No-Go stimulus (SI < 0, *n* = 124) were both significantly smaller than zero (\*\*\**P* < 2×10^-10^, Wilcoxon two-sided signed rank test). **n**, Modulation index for early and late response components for V1 neurons from trained mice. Early component, *P* = 0.06; late component, \*\*\**P* = 5.5×10^-47^, *n* = 394, Wilcoxon two-sided signed rank test. For **c**−**n**, source data are provided as a Source Data file. Shadings and error bars, mean ± s.e.m.

We performed single-unit extracellular recordings from V1 in behaving mice that had been trained for at least 5 days. We first compared the responses of V1 neurons to the Go and No-Go stimuli (Fig. 3e, f). We defined a selectivity index (SI) as (R_Go_ - R_No-Go_)/(R_Go_ + R_No-Go_), where R_Go_ and R_No-Go_ are firing rates to the Go and No-Go stimuli during the waiting period, respectively. The SIs of V1 neurons were significantly larger than zero (*P* = 2.2×10^-12^, Wilcoxon signed rank test) and significantly larger than those in untrained mice passively viewing the stimulus (*P* = 4.6×10^-4^, *n* = 394 neurons from trained mice and 160 neurons from untrained mice, Wilcoxon rank sum test, Fig. 3g, h), indicating a higher response to the Go stimulus than to the No-Go stimulus, consistent with previous reports^41^. While the SIs for the early response component (< 100 ms) were not significantly different from zero, the SIs for the late response component (>100 ms) were significantly larger than zero (Fig. 3i). We also compared the responses of V1 neurons to the No-Go stimulus between CR and FA trials (Fig. 3j, k). We defined a modulation index (MI) as (R_CR_ - R_FA_)/(R_CR_ + R_FA_), where R_CR_ and R_FA_ are firing rates to the No-Go stimulus during the waiting period in CR and FA trials, respectively. We found that the MIs were significantly smaller than zero for the population of V1 neurons (*P* = 1.46×10^-42^, *n* = 394 neurons, Wilcoxon signed rank test, Fig. 3l). For neurons preferring the Go stimulus (SI > 0, *n* = 270) and those preferring the No-Go stimulus (SI < 0, *n* = 124), the MIs were both significantly smaller than zero (Fig. 3m). Similar to the SI, the magnitude of MI was significant for the late but not for the early response component (Fig. 3n). In addition, we classified the neurons into broad-spiking and narrow-spiking cells^45,46^, which correspond to putative excitatory and putative inhibitory neurons, respectively. We found that the SIs of both cell types were significantly larger than zero, and the MIs of both cell types were significantly smaller than zero (Supplementary Fig. 4). These results suggest that, during the waiting period, the responses of V1 neurons to the reward-irrelevant No-Go stimulus were lower than those to the Go stimulus, and the responses to the No-Go stimulus were lower in trials in which the animals performed correctly than in trials performed incorrectly.

The stimulus selectivity and response modulation of V1 neurons during the waiting period could be attributed to multiple factors, including movement, reward expectation and top-down modulation^41,47^. We first analyzed whether V1 neurons exhibit licking-movement-related activity. We computed Pearson’s correlation coefficients between spike rate of V1 neurons and lick rate during the waiting period. The distribution of correlation was not significantly different from zero (Supplementary Fig. 5a). For the lick-triggered spike histogram computed using licks in the waiting period, we did not observe increase in V1 responses after the time of lick (Supplementary Fig. 5b). We also analyzed the responses during the waiting period using those trials in which no lick occurred within the first 0.5 s following stimulus onset. For these no-lick trials, the responses to the Go stimulus were significantly larger than those to the No-Go stimulus, and the responses to the No-Go stimulus remained significantly lower in CR than in FA trials (Supplementary Fig. 5c−f). Thus, the response modulation in V1 could not be attributed to licking movement.

We next examined whether the OFC top-down projection plays a role in the modulation of V1 responses in the Go/No-Go task. We expressed inhibitory opsin Jaws in the OFC, and recorded from V1 neurons with and without optogenetic inactivation of OFC axons in behaving mice. Laser stimulation was turned on 100-ms before or at stimulus onset, covering the duration of stimulus presentation, and laser-on and laser-off blocks were interleaved. For one group of mice, laser stimulation was applied during Go trials (Fig. 4a−d). We found that inactivating OFC axons in V1 during Go trials did not affect V1 responses to the Go stimulus during the waiting period (Fig. 4c), nor the behavioral performance of the mice (Fig. 4d). For another group of mice, laser stimulation was applied during No-Go trials (Fig. 4e−p). Inactivating OFC axons during No-Go trials significantly increased the responses of V1 neurons to the No-Go stimulus during the waiting period (*P* = 0.01, *n* = 169 neurons, Wilcoxon signed rank test), resulting in a significant reduction of SI (*P* = 0.007, Wilcoxon signed rank test, Fig. 4f). The increase of response to the No-Go stimulus was significant in CR but not in FA trials (CR: *P* = 1.02×10^-6^; FA: *P* = 0.66, *n* = 169 neurons, Wilcoxon signed rank test, Fig. 4h), leading to a reduction in the amplitude of MI (*P* = 6.4×10^-4^, Wilcoxon signed rank test, Fig. 4i). Such effect was observed in both experiments in which laser stimulation was turned on 100-ms before stimulus onset (Supplementary Fig. 6a−c) and at stimulus onset (Supplementary Fig. 6e−g). We also separately analyzed the effect of laser stimulation on the early and late response components. We found that inactivating OFC axons during No-Go trials did not significantly change the firing rates and MI for the early component, but significantly increased firing rates in CR trials and reduced the amplitude of MI for the late component (Fig. 4j−m). As laser stimulation during No-Go trials did not affect the mice’s lick rate or orofacial movement during the waiting period in CR trials (Fig. 4n, o, Supplementary Fig. 7 and Supplementary Video 1), the laser-induced response increase in CR trials could not be accounted for by a change in movement. Although inactivating OFC axons during No-Go trials significantly influenced the responses of V1 neurons to the No-Go stimulus, it did not change the performance of the mice (Fig. 4p). For control mice that GFP was expressed in the OFC, laser stimulation during No-Go trials did not significantly change the responses of V1 neurons to the No-Go stimulus in either CR or FA trials (Supplementary Fig. 6i, j). These data suggest that the OFC top-down projection to V1 contributes to the suppression of late-component V1 responses to the No-Go stimulus in CR trials.

**Fig. 4.**
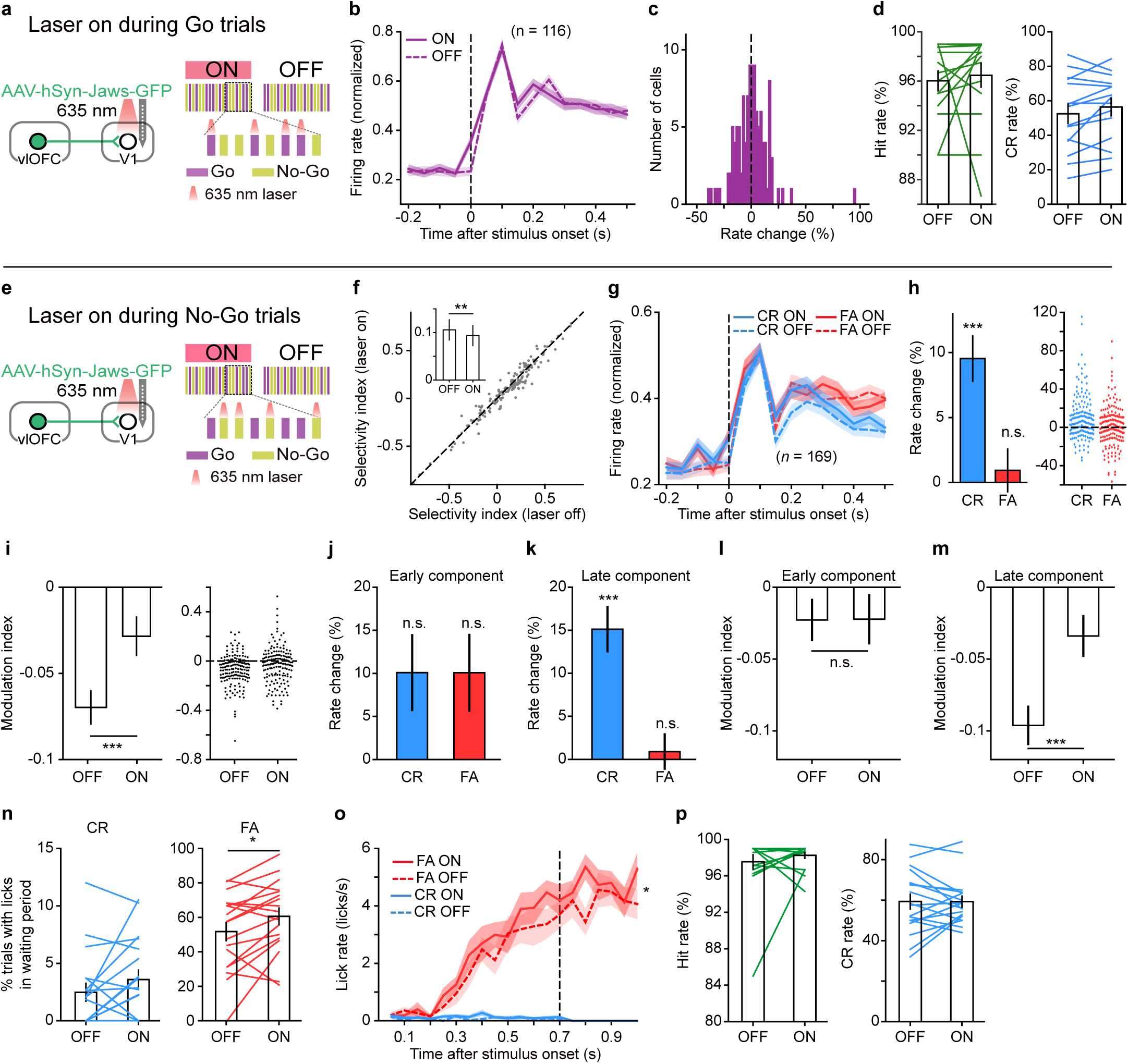
Optogenetic inactivation of OFC projection to V1 during No-Go trials increases V1 responses to No-Go stimulus in CR condition. **a**−**d**, Laser stimulation was applied during Go trials. **a**, Left, strategy for inactivating OFC projection to V1. Right, schematic of laser stimulation. **b**, Normalized responses to the Go stimulus with and without inactivating OFC axons in V1. **c**, Distribution of rate change induced by inactivating OFC axons in V1 during Go trials. *P* = 0.87, *n* = 116 neurons. The rate change was computed as (R_laser_on_ - R_laser_off_)/R_laser_off_, where R_laser_on_ and R_laser_off_ represented waiting-period firing rates to the Go stimulus with and without laser stimulation, respectively. **d**, Inactivating OFC axons in V1 during Go trials did not affect the hit rate (*P* = 0.26) or the CR rate of the mice (*P* = 0.11, *n* = 15 sessions from 7 mice). **e**−**p**, Laser stimulation was applied during No-Go trials. **e**, Strategy for inactivating OFC projection to V1 and schematic of laser stimulation. **f**, Inactivating OFC axons in V1 during No-Go trials significantly reduced the selectivity index of V1 neurons (\*\**P* = 0.007, *n* = 169). **g**, Normalized responses to the No-Go stimulus in FA and CR trials, respectively, with and without inactivating OFC axons in V1. **h**, Inactivating OFC axons in V1 during No-Go trials increased the responses of V1 neurons in CR (\*\*\**P* = 1.02×10^-6^) but not in FA trials (*P* = 0.66). *n* = 169 neurons. **i**, Inactivating OFC axons in V1 during No-Go trials significantly reduced the amplitude of MI (\*\*\**P* = 6.4×10^-4^, *n* = 169 neurons). **j**, Rate change for the early response component (CR, *P* = 0.18; FA, *P* = 0.77). **k**, Rate change for the late response component (CR, \*\*\**P* = 2.4×10^-5^; FA: *P* =0.75). **l**, Laser-off vs laser-on MI for the early response component (*P* = 0.78). **m**, Laser-off vs laser-on MI for the late response component (\*\*\**P* = 6.4×10^-5^). **n**, Inactivating OFC axons in V1 during No-Go trials increased the percentage of FA trials in which licks occurred within the waiting period (*P* = 0.02), but did not significantly change the percentage of CR trials in which licks occurred within the waiting period (*P* = 0.14, *n* = 18 sessions from 14 mice). **o**, Inactivating OFC axons in V1 during No-Go trials significantly increased the lick rate during waiting period of FA trials (two-way repeated measures ANOVA *F*_(1, 17)_ = 7.78, *P* = 0.01) but did not significantly change the lick rate during waiting period of CR trials (two-way repeated measures ANOVA *F*_(1, 17)_ = 1.6, *P* = 0.22). **p**, Inactivating OFC axons in V1 during No-Go trials did not affect the hit rate (*P* = 0.56) or the CR rate of the mice (*P* = 0.95, *n* = 18 sessions from 14 mice). Wilcoxon two-sided signed rank test was used for **c**, **d**, **f**, **h**−**n** and **p**. For **b**−**d** and **f**−**p**, source data are provided as a Source Data file. Shadings and error bars, mean ± s.e.m.

### Response patterns of V1-projecting OFC neurons

Next, we sought to identify V1-projecting OFC neurons with optogenetic tagging method and examined their responses during the Go/No-Go task. To this end, we first injected rAAV2-retro-hSyn-ChrimsonR-GFP^48^ in V1, resulting in the expression of excitatory opsin ChrimsonR in V1-projecting OFC neurons (Supplementary Fig. 8). We performed extracellular recordings from the OFC to monitor the spikes evoked by red laser stimulation (Fig. 5a, b). Using stimulus-associated spike latency test (SALT) for optogenetic identification^49^, we identified V1-projecting OFC neurons as those showing significant laser-evoked responses with short latencies (n = 22 out of 1175 units, *P* < 0.01, Fig. 5c, d and Supplementary Fig. 8f). In mice that had been trained for at least 6 days, the responses of identified OFC neurons to the Go and No-Go stimuli were monitored during the behavioral task (Fig. 5e). We defined a response index (RI) as (R_evoked_ - R_baseline_)/(R_evoked_ + R_baseline_), where R_evoked_ and R_baseline_ represented the firing rates during the waiting period and the baseline period before stimulus onset, respectively. For each neuron, we computed RIs for the responses to the Go stimulus in hit trials and those to the No-Go stimulus in CR (or FA) trials, respectively. We found that the RIs in hit and FA trials were both significantly smaller than zero, whereas those in CR trials were not significantly different from zero (Supplementary Fig. 8g−i). Thus, the responses to the Go stimulus in hit trials and those to the No-Go stimulus in FA trials were reduced as compared to the baseline responses, whereas those to the No-Go stimulus in CR trials were unchanged. During the behavioral sessions of OFC recordings, the mice showed anticipatory licking during the waiting period for the No-Go stimulus in FA trials as well as for the Go stimulus in hit trials (Supplementary Fig. 8j).

**Fig. 5.**
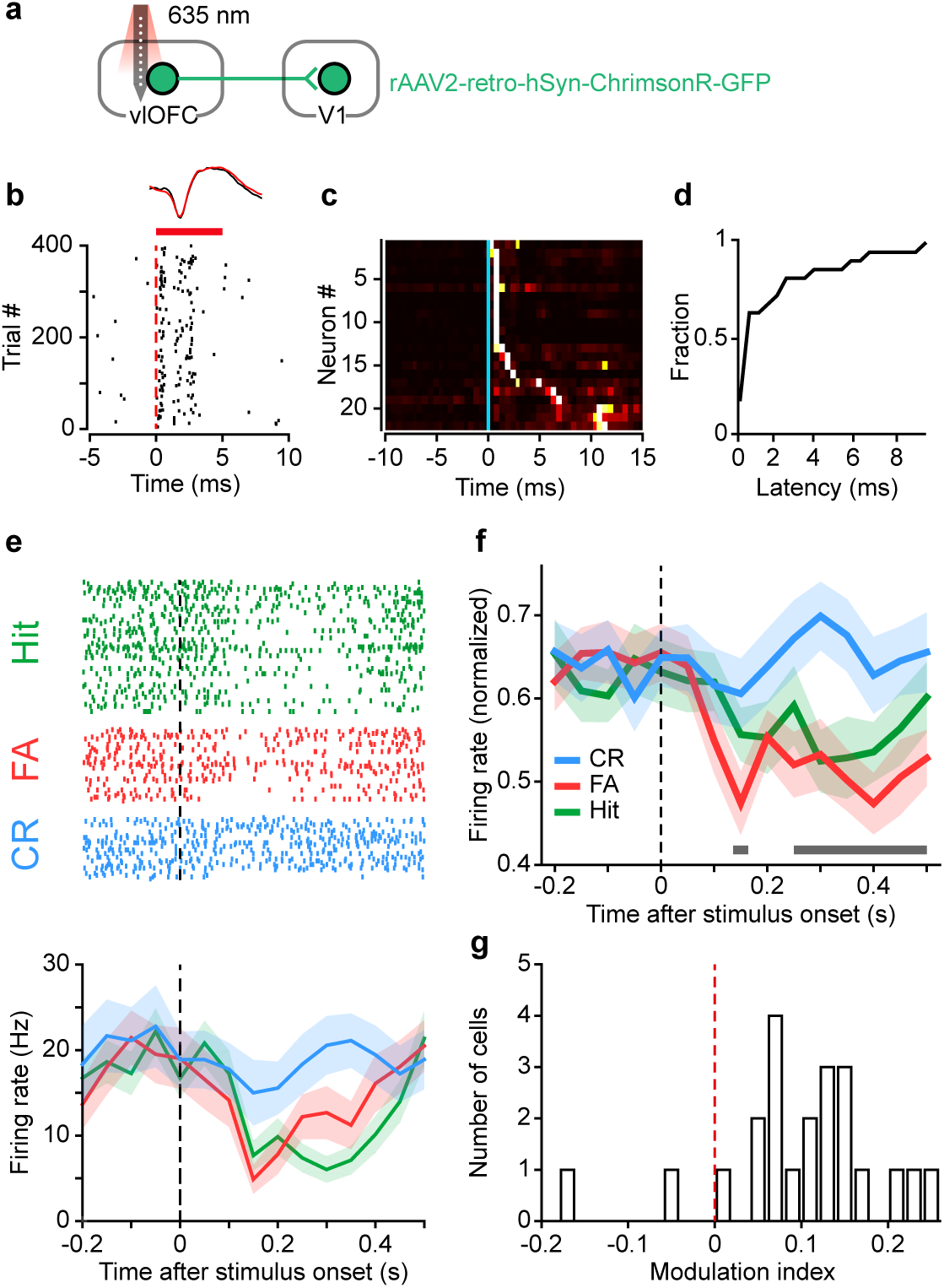
V1-projecting OFC neurons show response reduction during expectation of reward. **a**, Schematic of the strategy for phototagging V1-projecting OFC neurons. **b**, Spike raster of an identified V1-projecting OFC neuron aligned to laser onset. Red bar, duration of laser stimulation. Top, mean waveforms for spontaneous (black) and laser-evoked (red) spikes. **c**, PSTHs aligned to laser onset (blue line) for all identified neurons. The responses were normalized by peak value and sorted by peak latency. Colors from black to white indicate increase in firing rate. **d**, Cumulative distribution of laser-evoked spike latency for all identified neurons. **e**, Spike rasters and PSTHs of an identified V1-projecting OFC neuron during hit (green), FA (red) and CR (blue) trials, aligned to stimulus onset. **f**, The firing rates of each neuron were normalized by the maximum of the peak values in hit, FA, and CR trials, and were averaged across neurons. Horizontal bar indicates time points in which the responses between FA and CR conditions were significantly different (*P* < 0.05, two-way repeated measures ANOVA *F*_(1, 21)_ = 17.86, *P* = 3.79×10^-4^ followed by Sidak’s multiple comparisons test). **g**, Distribution of MIs for identified V1-projecting OFC neurons. *P* = 6.9×10^-4^, *n* = 22 neurons, Wilcoxon two-sided signed rank test. For **c**−**g**, source data are provided as a Source Data file. Shadings, mean ± s.e.m.

These results suggest that V1-projecting OFC neurons reduced firing when the mice expected reward. For V1-projecting OFC neurons, the responses to the No-Go stimulus were significantly lower in FA than in CR trials (Fig. 5f), and MIs were significantly larger than zero (*P* = 6.9×10^-4^, *n* = 22 neurons, Wilcoxon signed rank test, Fig. 5g), indicating that their responses exhibit outcome-expectancy modulation opposite to that found for V1 neurons. The spike rate of V1-projecting OFC neurons was not correlated with the lick rate (Supplementary Fig. 8k), and the histogram of lick-triggered spikes did not show increase in response following lick (Supplementary Fig. 8l, m). For those trials in which no lick occurred within the first 0.5 s following stimulus onset, the MIs of V1-projecting OFC neurons remained significantly larger than zero (Supplementary Fig. 8n). Thus, the response difference between CR and FA trials for these OFC neurons could not be attributed to licking. Given that activating OFC axons in V1 caused response suppression of V1 neurons (Fig. 1), the higher responses of V1-projecting OFC neurons in CR than in FA trials (Fig. 5f, g) may provide an explanation of the observation that V1 responses to the No-Go stimulus were lower in CR trials (Fig. 3k, l).

### OFC projection to V1 contributes to learning

The above results showed that, although inactivating OFC projection to V1 could affect V1 responses, it did not change the behavioral performance (Fig. 4p). As these mice had been trained before the optogenetic perturbation, we next examined whether perturbation of OFC projection from the first day of training influences the learning process. To inactivate the OFC projection to V1, we expressed Jaws in the OFC (Fig. 6a and Supplementary Fig. 9). For control mice, EGFP alone was expressed in the OFC (Fig. 6a). Both the Jaws-expressing and the EGFP-expressing mice were divided into two groups. The laser stimulation was applied to V1 during No-Go trials for one group (Fig. 6b and Supplementary Fig. 9) and during Go trials for another group (Fig. 6c and Supplementary Fig. 9). Throughout the learning process, each session consisted of interleaved blocks of laser-on and laser-off trials (20 trials/block).

**Fig. 6.**
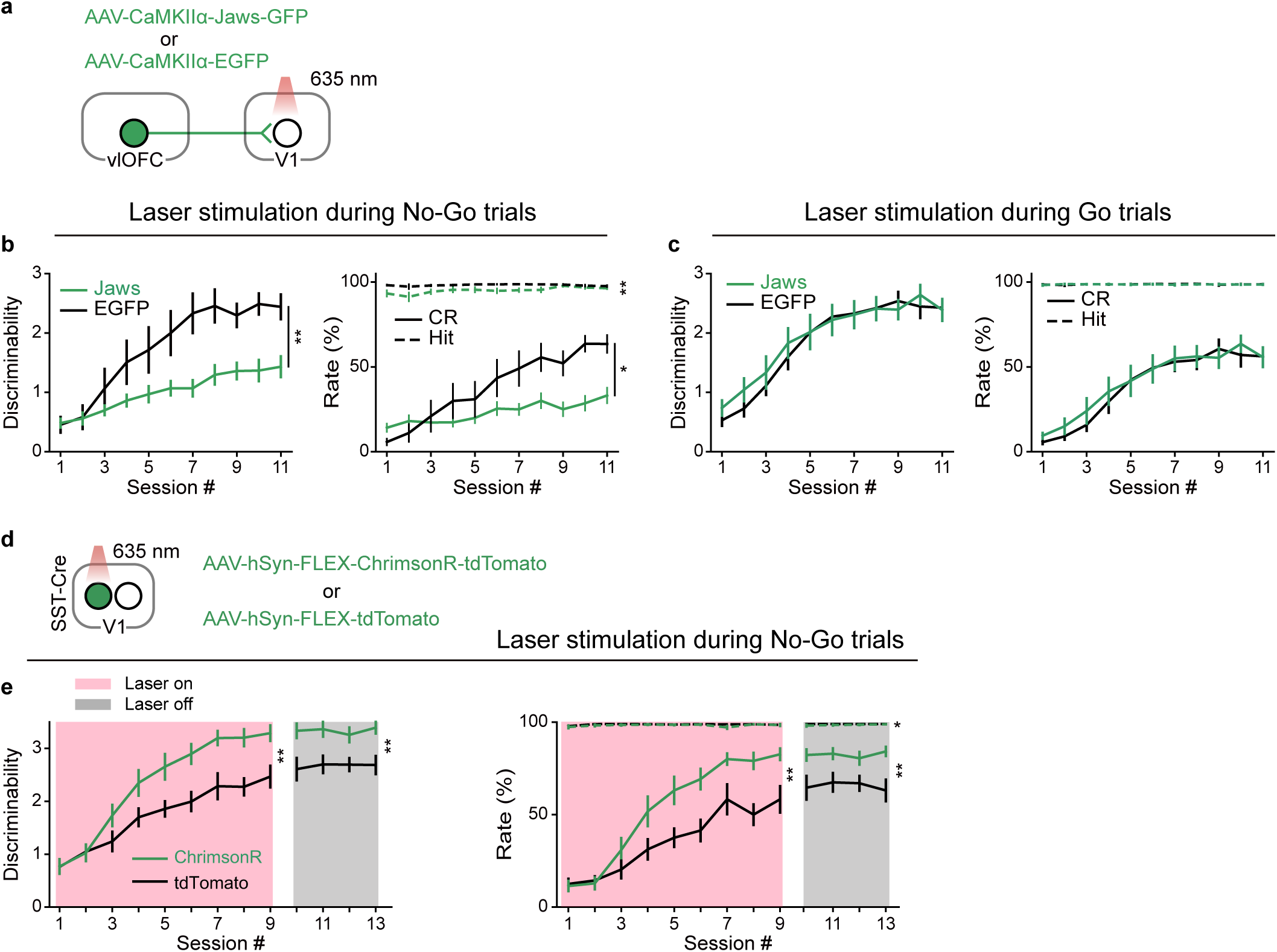
Effect of inactivating OFC projection to V1 or activating SST interneurons in V1 on the learning of Go/No-Go visual task. **a**, Schematic of the strategy for inactivating OFC projection to V1. **b**, Effect of inactivating OFC projection to V1 during No-Go trials. *n* = 9 and 13 for EGFP- and Jaws-expressing mice, respectively. For discriminability: *F*_(1, 20)_ = 10.24, \*\**P* = 0.004. For hit rate: *F*_(1, 20)_ = 13.38, \*\**P* = 0.002. For CR rate: *F*_(1, 20)_ = 6.69, \**P* = 0.02. Two-way ANOVA with mixed design. **c**, Effect of inactivating OFC projection to V1 during Go trials. *n* = 11 and 12 for EGFP- and Jaws-expressing mice, respectively. For discriminability: *F*_(1, 21)_ = 0.13, *P* = 0.72. For hit rate: *F*_(1, 21)_ = 0.24, *P* = 0.63. For CR rate: *F*_(1, 21)_ = 0.13, *P* = 0.72. Error bars, mean ± s.e.m. **d**, Schematic of the strategy for activating SST interneurons in V1. **e**, Effect of activating SST interneurons in V1 during No-Go trials. *n* = 10 and 11 for tdTomato- and ChrimsonR-expressing mice, respectively. Sessions 1**–**9, with laser stimulation; sessions 10–13, without laser stimulation. For discriminability in sessions 1**–**9: *F*_(1, 19)_ = 10.44, \*\**P* = 0.004; discriminability in sessions 10–13: *F*_(1, 19)_ = 11.93, \*\**P* = 0.003. For hit rate in sessions 1**–**9: *F*_(1, 19)_ = 4.37, *P* = 0.05; hit rate in sessions 10–13: *F*_(1, 19)_ = 6.07, \**P* = 0.02. For CR rate in sessions 1**–**9: *F*_(1, 19)_ = 10.63, \*\**P* = 0.004; CR rate in sessions 10–13: *F*_(1,19)_ = 9.84, \*\**P* = 0.005. For **b**, **c** and **e**, source data are provided as a Source Data file. Error bars, mean ± s.e.m.

For the group of Jaws-expressing mice with laser stimulation during No-Go trials, the behavioral performance were similar between laser-on and laser-off trials (Supplementary Fig. 9e), indicating that inactivation of the OFC axons in V1 during No-Go trials did not affect the immediate performance and corroborating the result in Fig. 4p. However, compared to the EGFP-expressing control mice, laser stimulation during No-Go trials slowed the learning in Jaws-expressing mice (Fig. 6b). For the experiments that laser stimulation was applied during Go trials, the performance was higher in laser-on than in laser-off trials for both the control and Jaws-expressing mice (Supplementary Fig. 9g, h), which may be due to the possibility that laser stimulation served as a cue to guide the mice’s behavior. Nevertheless, the Jaws-expressing and control mice did not differ in their learning curves (Fig. 6c), indicating that inactivating OFC axons in V1 during Go trials had no effect on learning. Together, the results demonstrate that learning to correctly reject the reward-irrelevant No-Go stimulus requires the activity of OFC projection to V1.

We further examined whether the learning process could be affected by optogenetic activation of the OFC top-down projection. We found that activation of OFC axons in V1 during No-Go or Go trials both caused an increase in behavioral performance in the session with laser stimulation, but the effect disappeared in the next session without laser stimulation (Supplementary Fig. 10). Such transient effect was likely due to antidromic activation of OFC neurons (Fig. 1g) that may activate other neurons or pathways involved in performing the task. Given that SST interneurons in V1 were innervated by OFC axons (Supplementary Fig. 2) and were the predominant interneuron subtype showing activity elevation following photostimulation of OFC neurons *in vivo* (Fig. 2i, j), we next examined the activity of SST interneurons in the Go/No-Go task and whether activating SST interneurons in V1 could affect learning. We used fiber photometry to measure the activity of SST interneurons by injecting AAV2/9-hSyn-FLEX-GCaMP6s-WPRE in V1 of SST-Cre mice (Supplementary Fig. 11a). In mice that had been trained for at least 4 days, the responses of SST interneurons to the No-Go stimulus during the waiting period were significantly higher in CR than in FA trials (Supplementary Fig. 11b, c). This response modulation was consistent with that found for the V1-projecting OFC neurons and may be partly attributed to the OFC projection to V1. To examine the role of SST interneurons in the learning of Go/No-Go task, we used SST-Cre mice in which AAVs encoding Cre-dependent ChrimsonR or tdTomato alone were injected into V1 (Fig. 6d). In each session of the learning process, laser stimulation of V1 was applied during No-Go trials, and blocks of laser-on and laser-off trials were interleaved. We found that activating SST interneurons in V1 during No-Go trials did not affect immediate performance in each session, as shown by similar d’ and CR rate between laser-on and laser-off trials (Supplementary Fig. 9j). However, the d’ and CR rate of ChrimsonR-expressing mice were significantly higher than those of control mice expressing tdTomato only (Fig. 6e).

After multiple sessions with laser stimulation during No-Go trials, the higher d’ and CR rate in ChrimsonR-expressing mice persisted on later sessions where laser stimulation was no longer applied (Fig. 6e), indicating that the improved performance was indeed due to learning. Thus, SST interneurons in V1 play an important role in the learning of correct rejection for the No-Go stimulus, serving as a substrate for modulating visual associative learning by the OFC to V1 projection.

Finally, we examined the relationship between behavioral learning rate and the degree to which OFC top-down projection modulates V1 responses. For another group of mice in which AAV-Jaws was injected in the OFC, behavioral training was conducted for 12 sessions without laser stimulation in V1. After the training, we performed extracellular recordings from V1, with and without laser stimulation in V1 during No-Go trials (Fig. 7a). For each mouse, we computed the laser-induced change in V1 firing rate to the No-Go stimulus in CR trials, and used the median of rate change of all neurons (22.9 ± 3.3 neurons/mouse, s.e.m.) to estimate the efficacy of OFC modulation of V1 responses. For each mouse, we also applied a reinforcement learning model^50,51^ to fit the curve of CR rate over the 12 sessions to obtain a learning rate (Fig. 7a, b). We found that the learning rate of the mouse correlated well with the laser-induced rate change of V1 neurons in CR trials (Fig. 7c). Thus, the data suggest that the individual differences in learning rate may be attributed to the efficacy of OFC top-down projection to modulate V1 responses.

**Fig. 7.**
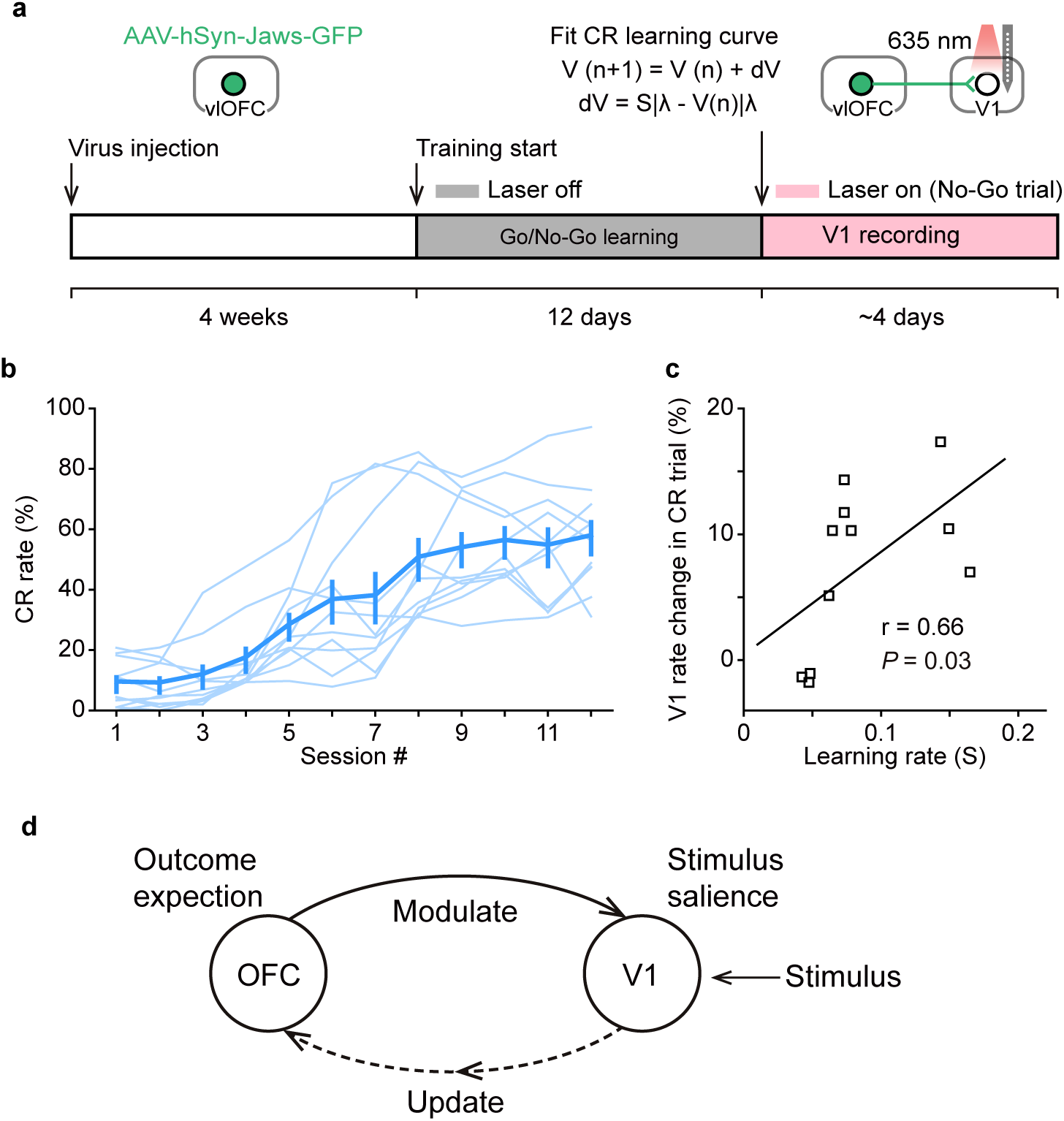
Behavioral learning rate correlates with the efficacy of OFC top-down projection to modulate V1 responses. **a**, Timeline of the experiments. **b**, CR rates over 12 training sessions for 11 mice. Each thin line represents a mouse. Error bars, mean ± s.e.m. The learning rate of each mouse was obtained by fitting the curve of CR rate with a reinforcement learning model. **c**, Laser-induced rate change of V1 neurons in CR trials correlated with learning rate (*r* = 0.66, *P* = 0.03). **d**, A schematic model for the role of OFC projection to V1 in associative learning. The OFC top-down projection mediates the outcome-expectancy modulation of V1 responses to reward-irrelevant stimulus. The response modification in V1 may represent a change in stimulus salience and may contribute to the update of outcome expectation signal in the OFC, potentially contributing to change of performance in future trials. For **b**, **c**, source data are provided as a Source Data file. Error bars, mean ± s.e.m.

## Discussion

In this study, we demonstrated that activation of OFC projection to V1 caused reduction of visual responses in V1 by preferentially recruiting SST interneurons in the local circuit. In mice performing a Go/No-Go visual task, the activity of OFC projection to V1 played an essential role in the outcome-expectancy modulation of V1 responses to the reward-irrelevant No-Go stimulus. Phototagging experiments showed that the responses of V1-projecting OFC neurons to the No-Go stimulus were decreased when the mice incorrectly expected reward. Importantly, we revealed that the OFC projection to V1 is critical for the learning of correctly rejecting the No-Go stimulus.

Top down projections to primary sensory areas play an important role in sensory processing and sensory-guided behavior^35,52^. Activating the projections from cingulate region of the frontal cortex to V1 enhances and suppresses V1 responses for the sites near and surround axonal activation, respectively, consistent with the effect of top-down modulation in selective attention^36^. Top-down projections from the anterior cingulate cortex to V1 carried stimulus prediction, which could be used to compute deviations from expectations and guide learning^39^. Associative learning also enhances the effect of top-down inputs from the retrosplenial cortex in modulating V1 responses^42^. In our study, we found that the responses of V1 neurons to the reward-irrelevant No-Go stimulus were modulated by outcome expectancy, being lower when the mice’ outcome expectation was correct than when it was wrong. Similar to that found in the somatosensory cortex^53,54^, the modulation of V1 responses by outcome expectation was evident in the late response component after stimulus onset. The response modulation in the late response component was reduced by optogenetic inactivation of OFC projection to V1, consistent with the role of OFC in the encoding of outcome expectation^2^. Chronic inactivation of OFC projection to V1 during No-Go trials slowed the improvement of CR rate, without affecting the performance per se, suggesting that the OFC top-down modulation of V1 responses to the No-Go stimulus is important for learning.

Few studies have recorded from behaving animals the responses of higher cortical neurons that provide top-down projections to V1^55^, and examined the causal role of top-down inputs to V1 in learning^56^. The CTB and virus retrograde tracing in our study showed that, the OFC neurons sending projections to V1 were in the ventrolateral OFC. The V1-projecting OFC neurons target all three subtypes of inhibitory interneurons in V1, similar to that found for OFC projections to auditory cortex^37^ and cingulate cortex projections to V1^36^. We found that optogenetic stimulation of OFC neurons *in vivo* preferentially activated SST interneurons in V1. To study the functional role of OFC projection to V1, we used a Go/No-Go visual task, in which the learning depended on the improvement of correct rejection for the reward-irrelevant No-Go stimulus. Many studies showed that the OFC neurons increased firing in anticipation of reward^2^, while some studies also found that a fraction of OFC neurons decreased firing to reward-predicting cues^27,31,57^. We found that, those OFC neurons not identified as V1-projecting showed higher firing rates to the No-Go stimulus in trials when mice incorrectly expected reward as well as to the Go stimulus when the reward expectation was correct (Supplementary Fig. 8o). By contrast, the V1-projecting OFC neurons exhibited response decrease relative to the baseline when the mouse expected reward for the Go stimulus in hit trials or for the No-Go stimulus in FA trials. This response decrease may account for the observation that inactivation of OFC projection to V1 during Go trials did not affect V1 responses to the Go stimulus or inactivation during No-Go trials did not affect responses to the No-Go stimulus in FA trials. For the No-Go stimulus, the responses of V1-projecting OFC neurons in CR trials were not different from the baseline response, and thus were higher than those in FA trials. Because the learning of Go/No-Go visual task manifested as an increase in the CR rate, the responses of V1-projecting OFC neurons to the No-Go stimulus likely increased with training as the percentage of CR trials increased. During No-Go trials, OFC projection to V1 may cause stronger activation of local SST interneurons in CR than in FA trials, leading to lower V1 responses in CR than in FA trials. Indeed, we found that the responses of SST interneurons in V1 to the No-Go stimulus were higher in CR than in FA trials, and inactivating OFC top-down projection increased V1 responses to the No-Go stimulus in CR trials. As the percentage of CR trials increased with training, the responses of V1 neurons to the No-Go stimulus likely decreased, representing a change in stimulus salience. Chronic activation of SST interneurons in V1 during No-Go trials facilitated learning, suggesting that recruitment of local SST interneurons by OFC top-down projection may promote learning by reducing V1 responses to reward-irrelevant stimulus. This result is also in line with recent reports that the activity of SST interneurons during learning may reflect the signals from long-range inputs^42,58^. Interestingly, we found that the individual differences in the learning rate may be partly attributed to the degree to which OFC top-down projection modulates V1 responses.

On the other hand, we found that optogenetic inactivation of OFC axons in V1 did not affect the immediate performance of mice. One possible explanation is that during the waiting period, the mice’s decision was determined by the responses of the OFC neurons, which could be seen from the divergence of response curves between FA and CR trials (Fig. 5f and Supplementary Fig. 8o), so that modulation of V1 responses would not affect the mice’s performance in current trial. A second possibility is that the effect of response modification in V1 needs to accumulate over multiple sessions to impact behavioral performance. As the OFC receives inputs from visual cortex^32-34^, the response modification in V1 may also in turn contribute to the update of outcome expectation signal in the OFC (Fig. 7d), potentially contributing to change of performance in future trials.

Studies using outcome devaluation and Pavlovian over-expectation tasks have revealed that the OFC is necessary for using expectations of specific outcome to guide behavior and learning^3,59^. The outcome predictions signaled by the OFC neurons could be utilized by downstream regions such as ventral striatum^60,61^, dorsal striatum^60,62^, BLA^63^ and VTA^23,29,64^. OFC lesion or inactivation disrupted the prediction errors signaled by dopaminergic neurons^29,64^ and expected reward values by putative non-dopaminergic neurons^64^ in VTA. While optogenetic inhibition of VTA-projecting OFC neurons did not impair the acquisition of Pavlovian trace conditioning, it impaired extinction learning and memory^27^. In addition, distinct OFC circuits were found to mediate different aspects of reinforcement learning in value-based decision making^65^. We found that the outcome expectation signals of OFC neurons are sent to V1 to regulate visual associative learning. As the OFC projects to other sensory cortices in addition to visual cortex^32^, the OFC top-down projection may also play an important role in associative learning for other sensory modalities.

## Methods

### Animals

All animal procedures were approved by the Animal Care and Use Committee at the Institute of Neuroscience, Chinese Academy of Sciences, and were in accordance with the guidelines of the Animal Advisory Committee at the Shanghai Institutes for Biological Sciences. We used the following mice: GAD67-GFP (CB6-Tg(Gad1-EGFP)G42Zjh), CaMKIIα-Cre (B6.Cg-Tg(Camk2a-cre)T29-1Stl), SST-Cre (Sst^tm2.1(cre)Zjh^), PV-Cre (B6;129P2-Pvalb^tm1(cre)Arbr^), VIP-Cre (Vip^tm1(cre)Zjh^), SST::Ai9 (generated by crossing SST-Cre with Ai9 mice, B6.Cg-Gt(ROSA)26Sor^tm9(CAG-tdTomato)Hze^), PV::Ai9, VIP::Ai9 and C57BL/6 mice. Adult (2 – 4 months) male mice were used for all experiments.

### Adeno-Associated Virus (AAVs)

We used the following AAVs: rAAV2-retro-hSyn-Cre (titer: 3.2 ×10^12^ viral particles/ml) and AAV2/8-EF1a-DIO-EYFP-WPRE (titer: 6.9×10^12^ viral particles/ml, for visualizing OFC axons in V1 in Fig. 1b); AAV2/8-CaMKIIα-hChR2(H134R)-mCherry (titer: 8.26×10^12^ viral particles/ml) and AAV2/8-hSyn-ChrimsonR-GFP (titer: 6.58×10^12^ viral particles/ml) (for activating OFC projection to V1 in Fig. 1, Supplementary Fig. 1 and Supplementary Fig. 10, or for activating OFC in Fig. 2g−k); AAV2/8-CaMKIIα-hChR2(H134R)-EYFP (titer: 8.66×10^12^ viral particles/ml; for activating OFC projection to V1 in Fig. 2a−f); AAV2/9-hSyn-FLEX-GCaMP6s-WPRE (titer: 6.9×10^12^ viral particles/ml; for fiber photometry experiment in Fig. 2g−k and Supplementary Fig. 11); AAV2/9-EF1α-DIO-His-EGFP-2A-TVA-WPRE (titer: 1.26×10^12^ viral particles/ml), AAV2/9-EF1α-DIO-RVG-WPRE (titer: 3.12×10^12^ viral particles/ml) and RV-EnvA-ΔG-dsRed (titer: 1×10^8^ viral particles/ml)(for retrograde monosynaptic tracing in Supplementary Fig. 2); AAV2/8-hSyn-Jaws-KGC-GFP-ER2 (titer: 5.4×10^12^ viral particles/ml; for inactivating OFC projection to V1 in Fig. 4, Fig. 7, Supplementary Fig. 1, Supplementary Fig. 6 and Supplementary Fig. 7); rAAV2-retro-hSyn-ChrimsonR-GFP (titer: 5.63×10^12^ viral particles/ml; for optogenetic tagging of V1-projecting OFC neurons in Fig. 5 and Supplementary Fig. 8); AAV2/8-CaMKIIα-Jaws-KGC-GFP-ER2 (titer: 5.38×10^12^ viral particles/ml; for inactivating OFC projection to V1 in Fig. 6 and Supplementary Fig. 9); AAV2/8-hSyn-FLEX-ChrimsonR-tdTomato (titer: 3.7×10^12^ viral particles/ml; for activating SST interneurons in Fig. 6 and Supplementary Fig. 9); AAV2/8-CaMKIIα-EGFP-WPRE (titer: 5.8×10^12^ viral particles/ml), AAV2/8-CaMKIIα-mCherry (titer: 5.7×10^12^ viral particles/ml), AAV2/8-hSyn-FLEX-tdTomato (titer: 5.1×10^12^ viral particles/ml) and AAV-hSyn-EGFP-WPRE (titer: 7.3×10^12^ viral particles/ml) (for optogenetic or fiber photometry experiments as a control group in Fig. 2i, Fig. 6, Supplementary Fig. 1, Supplementary Fig. 6 and Supplementary Fig. 9).

### Surgery

The mice were anesthetized with a mixture of midazolam (5 mg/kg), fentanyl (0.05 mg/kg) and medetomidine (0.5 mg/kg), and were head-fixed in a stereotaxic apparatus. For behavioral experiments without optogenetic manipulation, head plates were implanted before behavioral training. For *in vivo* recording and behavioral experiments with optogenetic manipulation, head plates were implanted after the virus injection. The virus was injected with a glass pipette (10 – 20 μm tip diameter) using a syringe pump (Harvard Apparatus). To observe the OFC axons in V1, we injected rAAV2-retro-hSyn-Cre (100 nl) in V1 (AP, −3.5 mm; ML, 2.4 mm; DV, 0.5 mm) and AAV2/8-EF1a-DIO-EYFP-WPRE (500 nl) in the OFC (AP, 2.7 mm; ML, 0.88 mm; DV, 1.8 mm). To manipulate the OFC to V1 projection in C57BL/6 mice, a craniotomy was made above the right OFC (AP, 2.7 mm; ML, 0.88 mm), and 500 nl of AAV (AAV2/8-hSyn-Jaws-KGC-GFP-ER2, AAV2/8-CaMKIIα-Jaws-KGC-GFP-ER2, AAV2/8-CaMKIIα-hChR2(H134R)-mCherry, AAV2/8-CaMKIIα-hChR2(H134R)-EYFP, AAV2/8-hSyn-ChrimsonR-GFP; or the control virus: AAV2/8-CaMKIIα-EGFP-WPRE or AAV2/8-CaMKIIα-mCherry) were injected into the cortex at a depth of 1.8 mm. A rectangular region on the skull above V1 (AP, −3.2 to −3.8 mm; ML, 2.0 to 2.8 mm) ipsilateral to the OFC injection site was marked by cutting and permanent red ink. The marked skull region above V1 was covered with tissue glue (Vetbond, 3M) until optogenetic manipulation or *in vivo* recording. For the experiments to block antidromic spiking of OFC neurons caused by laser stimulation of OFC axons in V1, a cannula (0.41 mm diameter) was implanted 500 μm above the virus injection site in the OFC. For fiber photometry recording of OFC stimulation-evoked activity in V1, AAV2/8-CaMKIIα-hChR2(H134R)-mCherry (600 nl) was injected to the right OFC at a depth of 1.8 mm and AAV2/9-hSyn-FLEX-GCaMP6s-WPRE (300 nl) was injected to the right V1 at a depth of 0.5 mm in SST-Cre, PV-Cre or VIP-Cre mice.

Following the virus injection, one optical fiber (200 μm diameter, NA 0.37) was inserted 100 μm above the injection site in the OFC and another one touching the dura of injection site in V1. For fiber photometry recording of SST interneurons in V1 in behaving mice, AAV2/9-hSyn-FLEX-GCaMP6s-WPRE (300 nl) was injected to the right V1 at a depth of 0.5 mm in SST-Cre mice, and optical fiber (200 μm diameter, NA 0.37) was inserted to touch the dura of V1. For phototagging of V1-projecting OFC neurons, a craniotomy was made above the right V1 (AP, −3.5 mm; ML, 2.4 mm), and rAAV2-retro-hSyn-ChrimsonR-GFP (300 nl) were injected into the cortex at a depth of 0.5 mm. To activate SST interneurons in V1 of SST-Cre mice, AAV2/8-hSyn-FLEX-ChrimsonR-tdTomato (300 nl) were injected into V1 at a depth of 0.5 mm, and the craniotomy was protected with a silicone elastomer (Kwik-Sil, WPI). After the virus injection, a stainless-steel headplate was fixed to the skull using dental cement. For mice used for Go/No-Go behavior with optogenetic manipulation of OFC axons in V1 or SST interneurons in V1, dental cement mixed with 50% carbon powder was used to cover the skull except the region above V1. The mice were injected with carprofen (5 mg/kg) subcutaneously after the surgery for 3 days, and were allowed to recover with food and water *ad libitum* for at least 7 days.

For infusion of tetrodotoxin (TTX) in the OFC, 1 μl of TTX (4 μM) was injected into the OFC through the implanted cannula 0.5 – 1 h before the *in vivo* recording.

Fluorescently conjugated Cholera toxin subunit B (CTB-555, 2 μg/μl, 300 nl, Invitrogen) was injected unilaterally into V1 (AP, −3.5 mm; ML, 2.4 mm) in C57BL/6 or GAD67-GFP mice. In some C57BL/6 mice, both CTB and rAAV2-retro-hSyn-ChrimsonR-GFP were injected in V1 to estimate the percentage of OFC neurons co-labeled by GFP and CTB. The histology experiments were performed 2 weeks after the injection.

Glycoprotein-deleted (ΔG) and EnvA-pseudotyped rabies virus (RV-EnvA-ΔG-dsRed) was used for retrograde monosynaptic tracing from different types of V1 neurons^43^. TVA receptor and rabies glycoprotein were expressed in Cre-positive neurons by co-injection of AAV2/9-EF1α-DIO-His-EGFP-2A-TVA-WPRE and AAV2/9-EF1α-DIO-RVG-WPRE (300 nl) in V1 in CaMKIIα-Cre, PV-Cre, SST-Cre and VIP-Cre mice. RV-EnvA-ΔG-dsRed (300 nl) was injected in the same site two weeks later. The histology experiments were performed 8 d after the RV injection.

### *In vivo* extracellular recording

Recordings with optogenetic stimulation were performed at least 3 weeks after the virus injection. For anesthetized experiments, mice were injected with chlorprothixene (3.2 mg/kg) subcutaneously and anesthetized with urethane (0.7 – 1.0 g/kg) intraperitoneally. The mouse was head-fixed in a stereotaxic frame and its body temperature was maintained at 37 ℃ through a heating blanket (FHC Inc.). A craniotomy (∼ 1 mm diameter) was made above V1 (AP, −3.5 mm; ML, 2.4 mm). For recordings in awake mice, the body of the mouse was restricted in a circular plastic tube and the headplate was fixed to a holder attached to the stereotaxic apparatus. While the animal was anesthetized with isoflurane (1 – 2%), a craniotomy (∼ 1 mm diameter) was made above V1 (AP, −3.5 mm; ML, 2.4 mm) or OFC (AP, 2.7 mm; ML 0.88 mm). The dura was removed, and the craniotomy was covered by ∼1% agarose dissolved in artificial cerebral spinal fluid (ACSF) and protected by a silicone elastomer (Kwik-Cast, WPI). The mouse was allowed to recover from the anesthesia in home cage for at least 1 hour. The recordings were made with multi-site silicon probes (A1×16-3mm-50-177 or A1×16-5mm-50-177, NeuroNexus Technologies; ASSY-77.2-64-6, Diagnostic Biochips, Inc.) mounted on a manipulator (MP-225, Sutter Instrument Company). For some recordings, the silicon probe was coated with DiO (Invitrogen) to allow *post hoc* recovery of penetration track. After finishing the recordings from awake mice, the electrode was retracted. The craniotomy was cleaned with ACSF, covered with ∼1% agarose and protected with a silicone elastomer (Kwik-Cast, WPI). After the experiments, the mouse was euthanized by an overdose of sodium pentobarbital.

The neural responses were amplified and filtered using a Cerebus 64-channel system (Blackrock microsystems). Local field potential signals were sampled at 2 kHz with a wide-band front-end filter (0.3 – 500 Hz). Spiking signals were sampled at 30 kHz. To detect the waveforms of spikes, we band-pass filtered the signals at 250 – 7500 Hz and set a threshold at 3.5 s.d. of the background noise. Spikes were sorted offline using the Offline Sorter (Plexon Inc.) based on cluster analysis of principle component amplitude. Spike clusters were considered to be single units if the percentage of spikes with interspike interval < 1 ms was lower than 0.3% and the *P* value for multivariate analysis of variance tests on clusters was less than 0.05.

### Slice preparation and recording

We used C57BL/6, SST::Ai9, PV::Ai9 and VIP::Ai9 mice for slice recordings. Mice that had been injected with AAV2/8-CaMKIIα-hChR2(H134R)-EYFP in the OFC were anesthetized with isoflurane and perfused with ice-cold cutting solution containing the following (in mM): sucrose 234, KCl 2.5, NaH_2_PO_4_ 1.25, MgSO_4_ 10, CaCl_2_ 0.5, NaHCO_3_ 26 and glucose 11 (300 – 305 mOsm). The mouse brain was dissected, and coronal slices (300 μm) were prepared using a vibratome (Leica VT1200S) in the ice-cold cutting solution. The prepared brain slices of V1 were incubated in ACSF containing the following (in mM): NaCl 126, KCl 2.5, NaH_2_PO_4_ 1.25, MgCl_2_ 2, CaCl_2_ 2, NaHCO_3_ 26 and glucose 10 (300 – 305 mOsm) for 30 – 45 min at 34°C, and then kept at room temperature. The cutting solution and ACSF were bubbled with 95% O_2_ and 5% CO_2_.

Whole-cell recordings of layer 2/3 V1 neurons in voltage-clamp mode were made at room temperatures (25 – 28°C) with a Multiclamp 700B amplifier and a Digidata 1440A (Molecular Devices). The electrodes were filled with a Cs-based low Cl^-^ internal solution containing the following (in mM): CsMeSO_3_ 130, MgCl_2_ 1, CaCl_2_ 1, HEPES 10, QX-314 2, EGTA 11, Mg-ATP 2, Na-GTP 0.3 (pH 7.3, 295 mOsm). Excitatory and inhibitory currents were recorded at −70 mV and 0 mV, respectively. Different types of V1 inhibitory neurons were identified by tdTomato-expressing neurons in SST::Ai9, PV::Ai9 and VIP::Ai9 mice. Pyramidal neurons were identified based on the morphology of tdTomato-negative cells and verified by staining of biocytin, which was included in the internal solution. Picrotoxin (50 μM, Tocris) and NBQX (10 μM, Tocris) were used to block GABA_A_ receptor and AMPA receptor mediated currents, respectively. Data were sampled at 10 or 20 kHz and analyzed with pCLAMP 10 (Molecular Devices). To determine whether the laser stimulation-evoked EPSC is significant, we computed the baseline by averaging the responses within 10 ms before laser onset, and compared the peak amplitude of EPSC with the baseline using Wilcoxon signed rank test.

### Fiber photometry

For fluorescence Ca^2+^ recordings, light from a 473-nm LED was reflected by a dichroic mirror (MD498, Thorlabs). The emission signals collected through the implanted optical fiber in V1 were filtered by a bandpass filter (MF525-39, Thorlabs) and detected by a photomultiplier tube (PMT, R3896, Hamamatsu). The light at the tip of the optical fiber was adjusted to 10 – 30 μW to minimize bleaching. An amplifier converted the output of the PMT to voltage signals, which were digitized using a data acquisition card (USB6009, National Instrument) at 200 Hz with custom-written programs.

### Visual stimulation

For *in vivo* recording experiments, visual stimuli were presented on a 17” LCD monitor (Dell P170S, mean luminance of 35 cd/m^2^, refresh rate 60 Hz) placed 9 cm away from the eye contralateral to the recording site, subtending 112.6° × 124.2° of visual space. Gamma correction was used to calibrate the monitor. The position of the monitor was adjusted such that the receptive fields (RFs) of the recorded neurons were at the center of the monitor. To locate the RFs of V1 neurons, we presented sparse noise stimuli over a black background, in which a white square (21° × 21°) was flashed for 33.3 ms on a 112.6° square grid in a pseudorandom sequence (100 repeats). To measure orientation tuning with and without inactivating OFC axons in V1, we presented drifting gratings (96° × 96°, spatial frequency = 0.03 cycles/deg, temporal frequency = 2 Hz, contrast = 100%) at 12 different directions (spaced at 30°) in a random sequence. Each stimulus was repeated 14 times for both laser-off and laser-on conditions. Each trial of the stimulus started with 1 s of gray screen, followed by 0.5 s of the first frame of grating and 2 s of the drifting grating.

For behavioral experiments, oriented gratings (90° × 90°, spatial frequency = 0.04 cycles/deg, contrast = 100%) were presented on a 17” LCD monitor (Dell E1713S, mean luminance 40 cd/m^2^, refresh rate 60 Hz) placed ∼10 cm away from the eye contralateral to the recording site or virus injection site. The Go and No-Go stimuli were vertically and horizontally oriented gratings, respectively. In each trial, the vertically (horizontally) oriented grating was static during the waiting period and then drifting rightward (upward) during the answer period. The Go and No-Go trials were randomly interleaved.

### Behavioral task

Mice were water-deprived for 2 days before the behavioral training. During behavioral experiments, the mouse was head-fixed and sat in an acrylic tube within a training box. Tongue licks were detected by the interruption of an infrared beam or a capacitance touch sensor, and the delivery of water was controlled by a peristaltic valve (Kamoer). The mice went through a habituation phase and a conditioning phase before learning the Go/No-Go task. For habituation (2 days), the mouse learned to lick from a custom-made lickspout to get water reward every 4 s. For conditioning (2 – 3 days), the mouse was trained to lick in response to a vertically oriented grating stimulus.

The grating was static for 0.7 s or 0.5 s (waiting period) and then drifting for 2.2 s or 2.4 s (answer period). If a lick was detected during the answer period, the mouse was rewarded with 5.5 μl of water. For Go/No-Go task, the grating stimulus in each trial was static during the waiting period and drifting during the answer period. For some groups of mice (Fig. 3, Fig. 4, Fig. 5, Fig. 7, Supplementary Fig. 3−8 and Supplementary Fig. 11), the durations of waiting period and answer period were 0.7 s and 2.2 s, respectively. For other groups of mice (Fig. 6, Supplementary Fig. 9 and Supplementary Fig. 10), the durations of waiting period and answer period were 0.5 s and 2.4 s, respectively. Licking during the waiting period was neither rewarded nor punished. For a Go stimulus, if a lick was detected during the answer period, the mouse was rewarded with 5.5 μl of water upon lick detection (hit). The mouse was neither rewarded nor punished for a miss (no lick during the answer period of Go stimulus), CR (no lick during the answer period of No-Go stimulus) or FA (lick during the answer period of No-Go stimulus). During the inter-trial interval (ITI), the screen was blank and licking was punished by a timeout period of 4 s. Licking during the 4-s timeout period triggered another 4-s timeout unless no lick was detected during the timeout period or the accumulated timeout exceeded 20 s. Each mouse performed the task for 1 h in each session.

In a subset of behavioral sessions, we recorded images of the facial area ipsilateral to the V1 recording site with a point grey camera (30 Hz frame rate) and a 780 nm longpass filter. Infrared LEDs (840 nm) were used to illuminate the face of the mouse.

### Optogenetic stimulation

Optical activation of ChR2 (ChrimsonR) was induced by blue (red) light. Optical silencing by Jaws activation was induced by red light. A blue laser (473 nm) or a red laser (635 nm) (Shanghai Laser & Optics Century Co.) was connected to an output optical fiber and the laser was controlled by a stimulus generator (Master 9, A.M.P.I.).

For measuring synaptic inputs from OFC axons to V1 neurons during slice recording experiments, blue light (1 ms duration) was delivered through a 40×0.8 NA water immersion lens at a power of 50 mW.

To manipulate the activity of OFC axons in V1 for *in vivo* recording or behavioral experiments, we used a zoom fiber collimator to focus the laser beam (∼600 μm diameter) on V1 or on the center of the marked skull region above V1. A shield was mounted on the mouse’s head to prevent leakage of laser light to the eyes or to the screen.

For *in vivo* extracellular recording of V1 neurons with optogenetic manipulation, trials with and without laser stimulation were interleaved. Laser stimulation covered the duration of stimulus presentation. For V1 recordings with laser stimulation during Go trials, the laser was turned on 100-ms before the onset of visual stimulus (Fig. 4a−d). For V1 recordings with laser stimulation during No-Go trials, the laser was turned on 100-ms before stimulus onset for some mice (Fig. 7 and Supplementary Fig. 6a−d) and at stimulus onset for other mice (Supplementary Fig. 6e−j). Laser was at a power of 5 mW at collimator output for blue laser and 10 mW for red laser.

For behavioral experiments in which the OFC to V1 projection was optogenetically inactivated or SST interneurons in V1 were activated, laser-off and laser-on blocks (20 trials/block) were interleaved in each session. In laser-on blocks, laser stimulation was applied during No-Go trials and Go trials in two separate groups of mice, respectively. During trials with laser stimulation, the laser was turned on 100-ms before stimulus onset, and turned off at stimulus offset. Laser was at a power of 10 or 15 mW at collimator output. For fiber photometry recording of OFC stimulation-induced V1 activity, 100 – 150 pulses of blue laser light (500 ms duration, interval at 10 s) were delivered to the optical fiber implanted in the OFC.

For phototagging of the V1-projecting OFC neurons, red laser light (15 mW) was applied on the surface of cortical area above the OFC ipsilateral to the virus injection site in V1. We delivered 400 light pulses (each 5-ms long) at 0.5 Hz.

### Histology

At least two weeks after the tracer/virus injection in V1 or virus injection in the OFC, the mouse was deeply anesthetized with sodium pentobarbital (120 mg/kg) and was perfused with 60 ml saline followed by 60 ml paraformaldehyde (PFA, 4%). Brains were collected, fixed in 4% PFA (4℃) overnight, and then transferred to 30% sucrose in phosphate-buffered saline (PBS) until equilibration. Brain sections (40 μm) were cut using a cryostat (Microm). The floating sections were incubated with Hoechst (2 μM, Thermo Fisher Scientific) in PBS for 10 min. The sections were rinsed in PBS for 10 min, mounted onto glass slides and coverslipped with VECTASHIELD Antifade Mounting Medium (Vector Laboratories, H-1000). Fluorescence images were taken with a Nikon A1 (Nikon Co. Ltd.) confocal microscope or the VS120 (Olympus). Images were analyzed with ImageJ (NIH, US).

### Data analysis

Analyses were performed in MATLAB. For the behavioral experiments, hit rate was computed as N_hits_/(N_hits_ + N_misses_), where N_hits_ and N_misses_ are the numbers of hit and miss trials, respectively. FA rate = N_FAs_/(N_FAs_ + N_CRs_), and CR rate = N_CRs_/(N_FAs_ + N_CRs_), in which N_FAs_ and N_CRs_ are the numbers of FA and CR trials, respectively. The behavioral discriminability was quantified as norminv(hit rate) – norminv(FA rate), in which norminv is the inverse of the cumulative normal function^66^. For behavioral experiments in which the OFC to V1 projection was optogenetically inactivated or SST interneurons in V1 were activated (Fig. 6), we computed hit rate, CR rate or discriminability using all trials (including laser-off and laser-on trials) in a session.

For each trial of Go/No-Go task, we computed lick latency as the time of first lick within 1 s after stimulus onset. The lick latency in each session was quantified as the median of first lick latency across trials. For behaving mice used for V1 or OFC recordings, the lick latency (V1 recording: 476.3 ± 17.5 ms, s.e.m.; OFC recording: 447.2 ± 18.3 ms, s.e.m.) was close to 0.5 s even if the waiting period was 0.7 s. For the following analysis of neuronal responses during the waiting period, we thus used the responses within the first 0.5 s of the waiting period. To determine whether a V1 neuron was responsive to visual stimulus, we computed the baseline response during the 0.2 s before stimulus onset and the evoked firing rate during the waiting period of stimulus presentation for each trial. Those V1 neurons in which the evoked responses during the waiting period were larger than 0.5 spike/s and were significantly higher than the baseline response (*P* < 0.05, Wilcoxon signed rank test) were used in the analysis. For the responses to No-Go stimulus during the waiting period, we divided the trials into CR and FA conditions, and computed a modulation index (MI) as (R_CR_ - R_FA_)/(R_CR_ + R_FA_), in which R_CR_ and R_FA_ represented responses for CR and FA trials, respectively. For the analysis of MI, we only included neurons from those sessions in which the number of FA trials > 15. Statistical significance of the MI of each cell was determined by comparing the waiting-period responses to No-Go stimulus between CR and FA trials with Wilcoxon rank sum test. The selectivity of V1 responses to the Go and No-Go stimuli was evaluated by a selectivity index (SI), defined as (R_Go_ - R_No-Go_)/(R_Go_ + R_No-Go_), in which R_Go_ and R_No-Go_ are firing rates to the Go and No-Go stimuli during the waiting period, respectively.

For the experiments with optogenetic inactivation of OFC projection to V1 in behaving mice, we estimated the rate change of V1 neurons induced by laser stimulation in Go trials or No-Go trials. For Go trials, the rate change was computed as (R_laser_on_ - R_laser_off_)/R_laser_off_, where R_laser_on_ and R_laser_off_ represented waiting-period firing rates to Go stimulus with and without laser stimulation, respectively. For CR condition in No-Go trials, the rate change was computed as (R_CR_laser_on_ - R_CR_laser_off_)/R_CR_laser_off_, where R_CR_laser_on_ and R_CR_laser_off_ represented waiting-period firing rates to No-Go stimulus in CR trials with and without laser stimulation, respectively. For FA condition in No-Go trials, the rate change was computed as (R_FA_laser_on_ - R_FA_laser_off_)/R_FA_laser_off_, where R_FA_laser_on_ and R_FA_laser_off_ represented waiting-period firing rates to No-Go stimulus in FA trials with and without laser stimulation, respectively. Because some mice had few FA trials due to a high behavioral performance, we only included V1 neurons from those sessions in which the numbers of FA trials in laser-on and laser-off conditions were both > 15.

For extracellular recordings of V1 neurons from anesthetized or awake mice not performing behavioral task, we first estimated the RFs of neurons by cross-correlating the responses with the sparse noise stimuli^67^. For the responses to oriented drifting gratings, spike rate to each stimulus was calculated by averaging the responses during the drifting period over all trials. For the responses without laser stimulation, we calculated the t statistic (mean evoked rate divided by s.e.m.) for the responses to the preferred orientation^68^. For the responses in laser-off and laser-on conditions, respectively, we computed a global measure of orientation selectivity index (OSI) as:

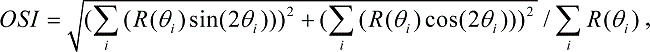

where *θ* _i_ is the angle of the drifting direction of the grating and *R* (*θ* _i_) is the response at angle *θ* _i_. Only those units with OSI > 0.08 (sensitive to orientation^69^), t > 2 (visually responsive^68^) and peak evoked firing rate > 2 Hz during laser-off condition were included in the subsequent analyses. To estimate the effect of activating OFC axons in V1 on the response amplitude of V1 neurons, we computed a rate change index as (R_laser_on_ - R_laser_off_)/(R_laser_on_ + R_laser_off_), in which R_laser_on_ and R_laser_off_ represented responses averaged over all orientations for laser-on and laser-off trials, respectively.

We divided the responses of V1 neurons during the waiting period into an early component (< 100 ms) and a late component (>100 ms). The SI and MI of each V1 neuron were computed for the early and the late components, respectively. Based on the spike waveforms, we classified V1 neurons as broad-spiking and narrow-spiking cells, which correspond to putative excitatory and putative inhibitory neurons, respectively^45,46^. For each cell, the spikes were aligned by the trough and averaged, and the average waveform was interpolated^46^. We computed peak width as the width of the peak at half-maximum of the peak amplitude. Based on the distribution of the peak width (Supplementary Fig. 4), we defined a threshold at 0.35 ms to classify broad-spiking and narrow-spiking cells. The SI and MI were computed for broad-spiking and narrow-spiking cells, respectively.

To identify V1-projecting OFC neurons, we used the stimulus-associated spike latency test (SALT)^49^ to determine whether laser stimulation significantly changed the spike timing of neurons after stimulation onset. We also calculated Pearson’s correlation coefficient between waveforms of spontaneous spikes and spikes during the 10-ms period after laser onset. A unit was identified as ChrimsonR-expressing neuron if *P* < 0.01 for SALT test and waveform correlation coefficient > 0.9. To determine the response latency relative to laser onset, we binned the spikes at 0.1-ms resolution. For the peri-stimulus time histogram (PSTH) within the 10-ms period after laser onset, we identified the time of peak firing rate. For each bin of laser-evoked response within the time of peak response, we tested the difference between the firing rate in this bin and that averaged over 10-ms duration before laser onset (t test)^70^. The latency was identified as the first time point after laser onset with *P* < 0.01.

For OFC neurons recorded from behaving mice, we computed the baseline firing rates during the 0.2 s before stimulus onset and the responses to visual stimuli during the waiting period. Those neurons in which the waiting-period responses were significantly different from the baseline responses (*P* < 0.05, Wilcoxon signed rank test) were used in the analysis. We computed MI for V1-projecting OFC neurons using the same equation for V1 neurons. For OFC neurons, we also computed a response index (RI) for the responses to Go stimulus in hit trials and the responses to No-Go stimulus in CR (or FA) trials, respectively. RI was defined as (R_evoked_ - R_baseline_)/(R_evoked_ + R_baseline_), where R_evoked_ and R_baseline_ represented the waiting-period firing rate and the baseline rate, respectively.

We performed several analyses to examine whether spike rate of V1 neurons or V1-projecting OFC neurons during the waiting period is affected by licking. First, we calculated the Pearson’s correlation coefficient between the spike rate and lick rate in FA trials. For V1 neurons (V1-projecting OFC neurons), those sessions with >20 (>5) FA trials in which licks occurred in the waiting period were used for this analysis. Correlation with *P* < 0.05 was considered as significant. Second, we used licks in the waiting period to compute histogram of lick-triggered spikes. For V1-projecting OFC neurons, we further divided the neurons into two groups, in which the firing rates during the waiting period in hit trials were lower (RI < 0) and higher (RI > 0) than the baseline rate, respectively.

Histogram of lick-triggered spikes were computed for both groups of V1-projecting OFC neurons. Third, we analyzed the selectivity index and modulation index of V1 neurons (V1-projecting OFC neurons) using those trials in which no lick occurred within the first 0.5 s following stimulus onset.

For fiber photometry experiments to measure V1 activity induced by OFC stimulation, the value of fluorescence change (ΔF/F) was derived by calculating (F - F_0_)/F_0_, where F_0_ is baseline fluorescence signal averaged over 2 s before laser stimulation. For each mouse, we computed the peak of ΔF/F values after laser onset and the latency of peak ΔF/F. For fiber photometry experiments to measure the activity of SST interneurons from behaving mice, F_0_ was computed using the signal averaged over 1 s before stimulus onset. To compute the responses to the No-Go stimulus in CR (or FA) trials, we averaged the values of ΔF/F during the waiting period.

The orofacial movements of the mice were analyzed using the FaceMap software (www.github.com/MouseLand/FaceMap)47. For each mouse, the motion energy PCs during the waiting period between laser-off and laser-on conditions were compared.

### Reinforcement learning model

We used a reinforcement learning model^50,51^ to fit the CR rate for the No-Go stimulus across sessions. For each session, the learned association (V) between the No-Go stimulus and the non-reward state was quantified by the CR rate. We assumed that the adjustment of V based on association prediction error was controlled by the subjective salience (S) of the No-Go stimulus. The association was updated according to the following equations:

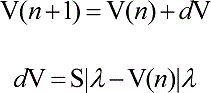

where λ is the intensity of unconditioned stimulus and set to 1, and *n* is the session number. We fitted the free parameter S by minimizing the sum of the squares of the residuals (least squares) using MATLAB (Mathworks). The S was used to quantify the learning rate of each mouse.

### Statistical analysis

Sample sizes were similar to others used in the field. No statistical method was used to predetermine sample size. The statistical analysis was performed using MATLAB or GraphPad Prism (GraphPad Software). Wilcoxon signed rank test (one-sided or two-sided), Wilcoxon two-sided rank sum test, one-way repeated measures ANOVA, two-way repeated measures ANOVA, or two-way ANOVA with mixed designed was used to determine the significance of the effect. Correlation values were computed using Pearson’s correlation. Data were not collected in a blinded fashion. Unless otherwise stated, data were reported as mean ± s.e.m.

## Data availability

Source data are provided as a Source Data file. The other data that support the findings of this study are available from the corresponding author upon reasonable request.

## Code availability

The data acquisition and analysis code are available from the corresponding author upon reasonable request.

## Supporting information

OFC to V1 paper 2020-3-21 for bioRxiv.pdf

## Acknowledgements

We thank M. M. Poo for comments on the manuscript. We thank Y. Li and W. Xu for technical assistance and Y.-N. Dou for help with slice recordings. This work was supported by the Strategic Priority Research Program of Chinese Academy of Sciences (grant No. XDB32010200), Shanghai Municipal Science and Technology Major Project (grant No. 2018SHZDZX05), and the National Natural Science Foundation of China (31571079, 31771151).

## Author contributions

D.L. and H.Y. conceived and designed the experiments. D.L. performed and organized all the experiments. J.D. and Y.-G.S. performed whole-cell recording experiments. Z. Z. and T.Y. performed reinforcement model fitting. D.L., Z.-Y. Z. and H.Y. analyzed the data. H.Y. wrote the manuscript with inputs from all authors. All authors read and revised the manuscript.

