## Supplementary material for "Orbitofrontal control of visual cortex gain promotes visual associative learning": OFC to V1 paper 2020-3-21 for bioRxiv.pdf

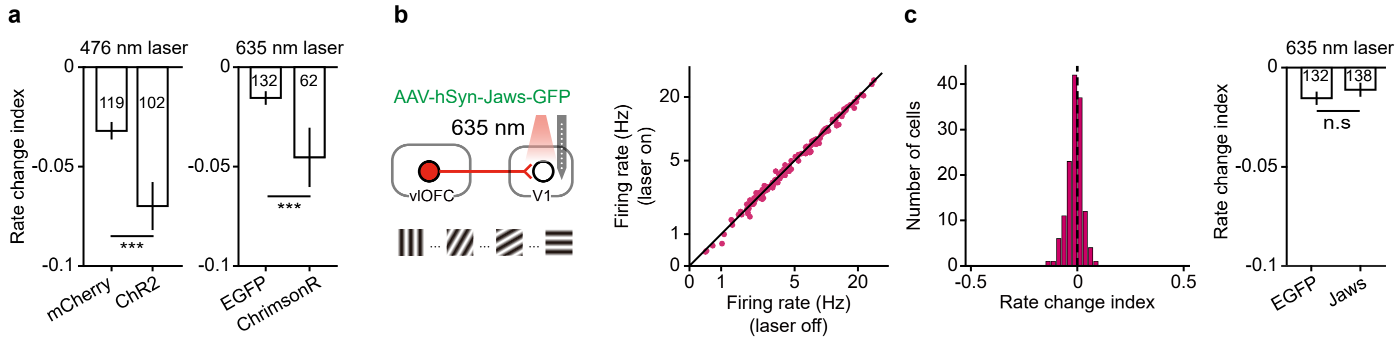


**Supplementary Figure 1 | Activating OFC projection to V1 reduces response amplitude of V1 neurons, whereas inactivating OFC projection to V1 does not affect V1 response amplitude in passive-viewing mice.**

**a**, Left, comparison of rate change index of V1 neurons between mice injected with AAV-mCherry and AAV-ChR2 in the OFC. The rate change was induced by blue laser stimulation in V1. The magnitude of rate change index was significantly higher for V1 neurons in ChR2-expressing mice ($n$ = 102 neurons) than in mCherry-expressing mice ($n$ = 119 neurons). ***$P$ = $5.9\times10^{-4}$. Right, comparison of rate change index of V1 neurons between mice injected with AAV-EGFP and AAV-ChrimsonR in the OFC. The rate change was induced by red laser stimulation in V1. The magnitude of rate change index was significantly higher for V1 neurons in ChrimsonR-expressing mice ($n$ = 62 neurons) than in EGFP-expressing mice ($n$ = 132 neurons). ***$P$ = $3.9\times10^{-4}$. Wilcoxon two-sided rank sum test. **b**, Left, schematic of measuring V1 visual responses with and without inactivating OFC axons in V1. Right, mean firing rate (firing rate averaged over all orientations) of V1 neurons with laser on vs. laser off. $P$ = $7.4\times10^{-4}$, $n$ = 138 neurons from awake, passive-viewing mice. **c**, Left, distribution of rate change indexes for V1 neurons shown in **b**. Right, the magnitude of rate change index was not significantly different between V1 neurons in EGFP-expressing mice ($n$ = 132 neurons) and Jaws-expressing mice ($n$ = 138 neurons). $P$ = 0.17, Wilcoxon two-sided rank sum test. Error bars, mean ± s.e.m. Source data are provided as a Source Data file.

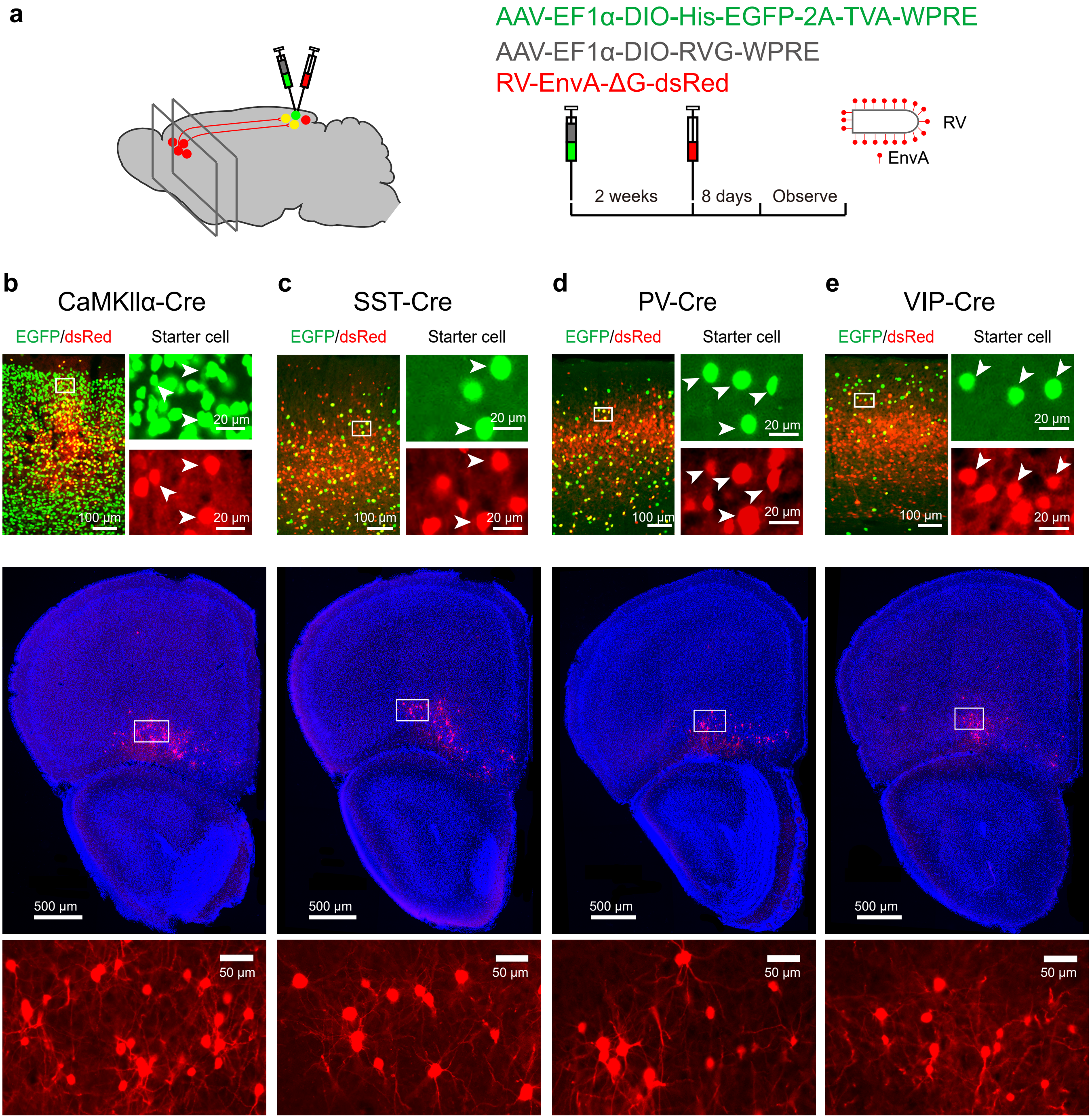


**Supplementary Figure 2 | Rabies virus-mediated trans-synaptic retrograde tracing reveals monosynaptic innervation of different cell types in V1 by OFC neurons.**
**a**, Viral injection procedure. **b**, Upper panel, injection site in V1 in a CaMKIIα-Cre mouse. Enlarged views of region in white box are shown on the right. Yellow or arrow heads, starter cells. Middle panel, fluorescence image showing trans-synaptically labeled dsRed-expressing neurons in the OFC. Enlarged view of region in white box is shown on the bottom. **c-e**, Similar to those described in **b** except that the images were from SST-Cre, PV-Cre and VIP-Cre mice, respectively.

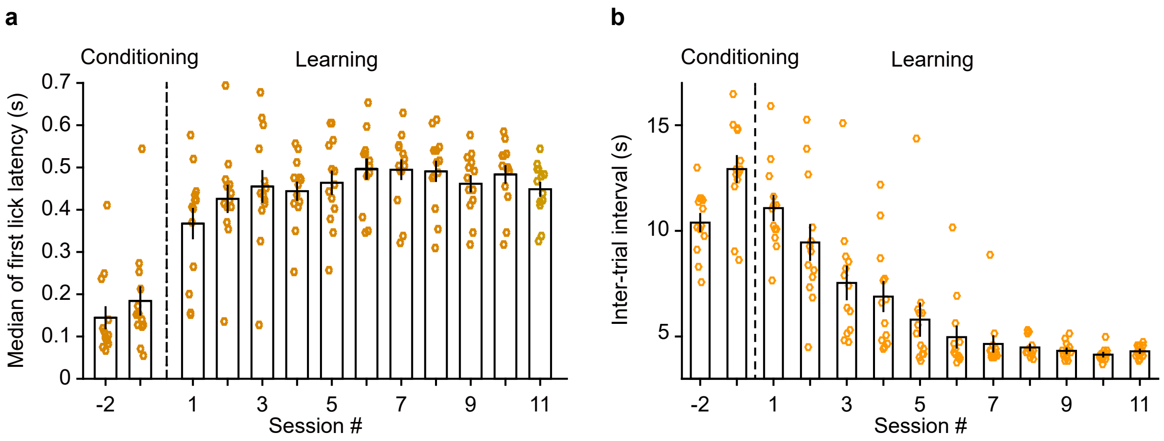


**Supplementary Figure 3 | Lick latency and inter-trial interval for mice across training sessions.**
**a**, Lick latency across sessions (2 sessions of conditioning and 11 sessions of learning the Go/No-Go task) for a population of mice ($F_{(3.03,\ 36.32)} = 21.81$, $P = 2.8\times10^{-8}$, $n = 13$). In each session, each circle represents the median lick latency across trials for a mouse. For each trial, lick latency was computed as the time of first lick after stimulus onset. **b**, Inter-trial interval across sessions (2 sessions of conditioning and 11 sessions of learning the Go/No-Go task) for a population of mice ($F_{(2.78,\ 33.37)} = 50.23$, $P = 3.76\times10^{-12}$). In each session, each circle represents the mean ITI for a mouse. Source data are provided as a Source Data file. Error bars, mean ± s.e.m.

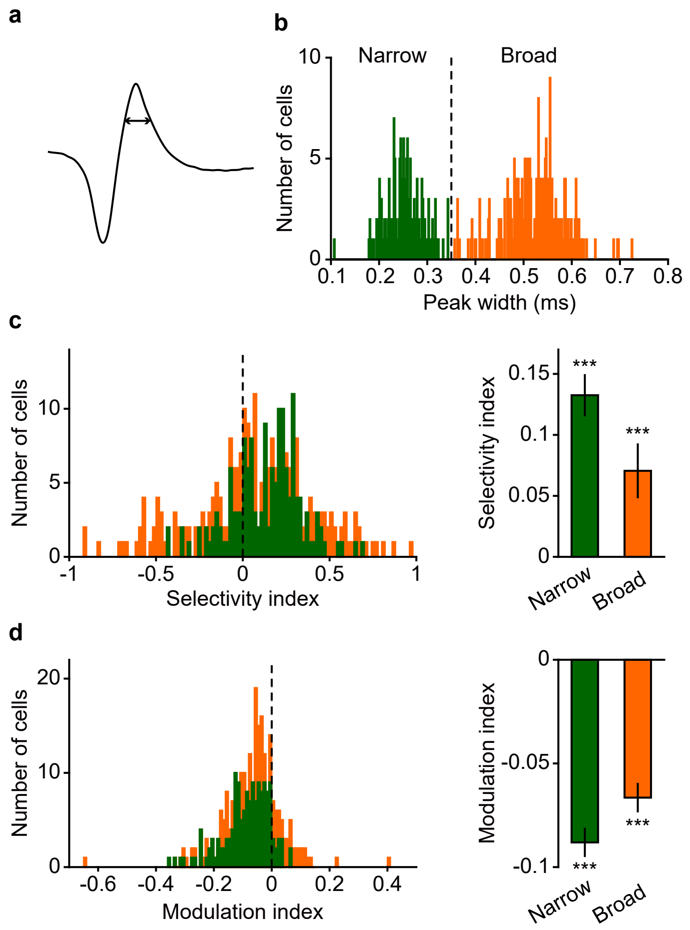


**Supplementary Figure 4 | Selectivity index and modulation index for broad-spiking and narrow-spiking V1 cells.** **a**, Average spike waveform for an example V1 neuron. Peak width was computed as the width of the peak at half-maximum of the peak amplitude. **b,** Distribution of peak width. A threshold at 0.35 ms (vertical dashed line) was used to segregate narrow-spiking cells (green, $n = 150$) from broad-spiking cells (orange, $n = 244$). **c**, The selectivity index was significantly larger than zero for both narrow-spiking cells (***$P = 9.6\times10^{-13}$) and broad-spiking cells (***$P = 2.7\times10^{-4}$). **d**, The modulation index was significantly smaller than zero for both narrow-spiking cells (***$P = 3.7\times10^{-23}$) and broad-spiking cells (***$P = 1.2\times10^{-21}$). Data were analyzed using Wilcoxon two-sided signed rank test. For **b-d**, source data are provided as a Source Data file. Error bar denotes s.e.m.

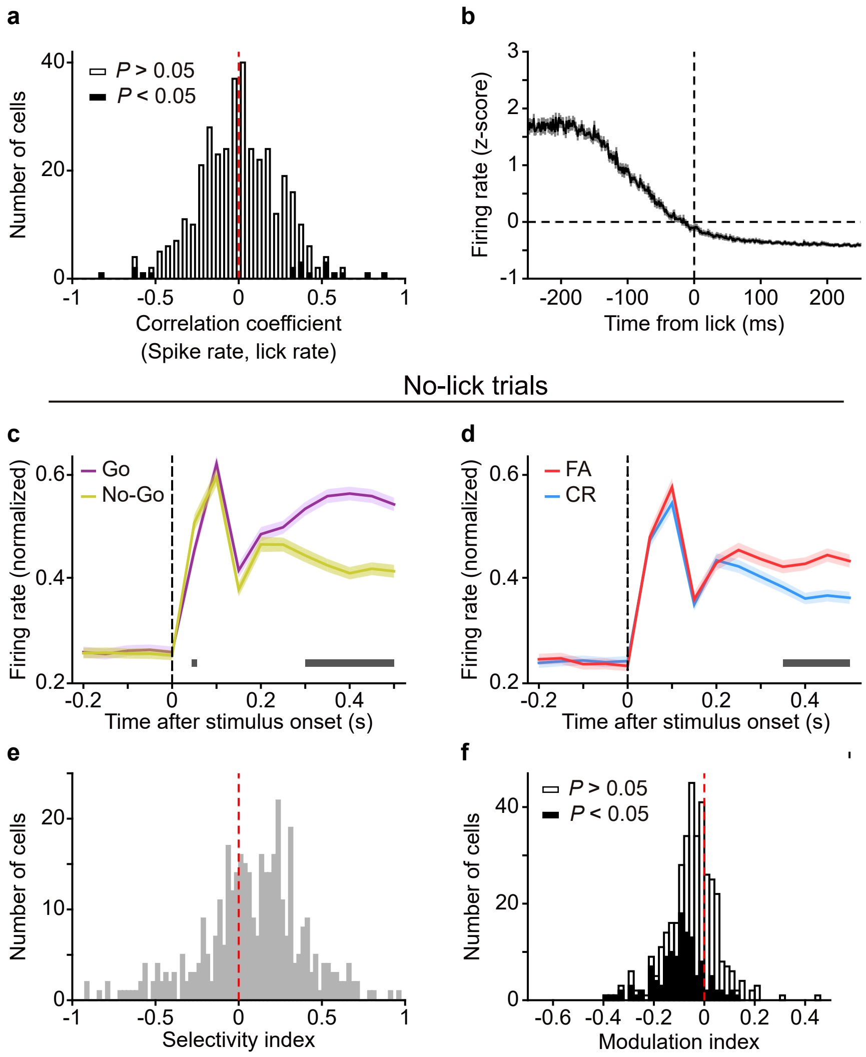


**Supplementary Figure 5 | Response modulation in V1 could not be attributed to licking movement.**
**a**, Distribution of Pearson's correlation coefficients between spike rate of V1 neurons and lick rate in FA trials during the waiting period ($P = 0.85$, $n = 364$). We only used those cells with > 20 FA trials in which lick occurred within the waiting period. Open and black bars denote cells with non-significant and significant coefficients, respectively. **b**, Histogram of lick-triggered spikes, computed using spikes and licks during the waiting period ($n = 364$ neurons). **c**, The responses of V1 neurons to the Go and No-Go stimuli after excluding those trials where lick occurred during the waiting period. The firing rates of each V1 neuron were normalized by the maximum of the peak values in Go and No-Go trials, and were averaged across neurons. Horizontal bar indicates time points in which the responses between the Go and No-Go stimuli were significantly different ($P < 0.05$, two-way repeated measures ANOVA $F_{(1, 376)} = 32.5$, $P = 2.4\times10^{-8}$ followed by Sidak's multiple comparisons test). **d**, The responses of V1 neurons to the No-Go stimulus in FA and CR trials after excluding those trials where lick occurred during the waiting period. Horizontal bar indicates time points in which the responses between FA and CR trials were significantly different ($P < 0.05$, two-way repeated measures ANOVA $F_{(1, 376)} = 71.8$, $P = 5.6\times10^{-6}$ followed by Sidak's multiple comparisons test). **e**, After excluding those trials in which lick occurred during the waiting period, the selectivity index of V1 neurons was still significantly larger than zero ($P = 1.0\times10^{-9}$, $n = 377$). **f**, After excluding those trials in which lick occurred during the waiting period, the modulation index of V1 neurons was still significantly smaller than zero ($P = 5.5\times10^{-20}$, $n = 377$). Open and black bars denote cells with non-significant and significant MIs, respectively. Wilcoxon two-sided signed rank test was used for **a**, **e** and **f**. Source data are provided as a Source Data file. Shadings denote s.e.m.

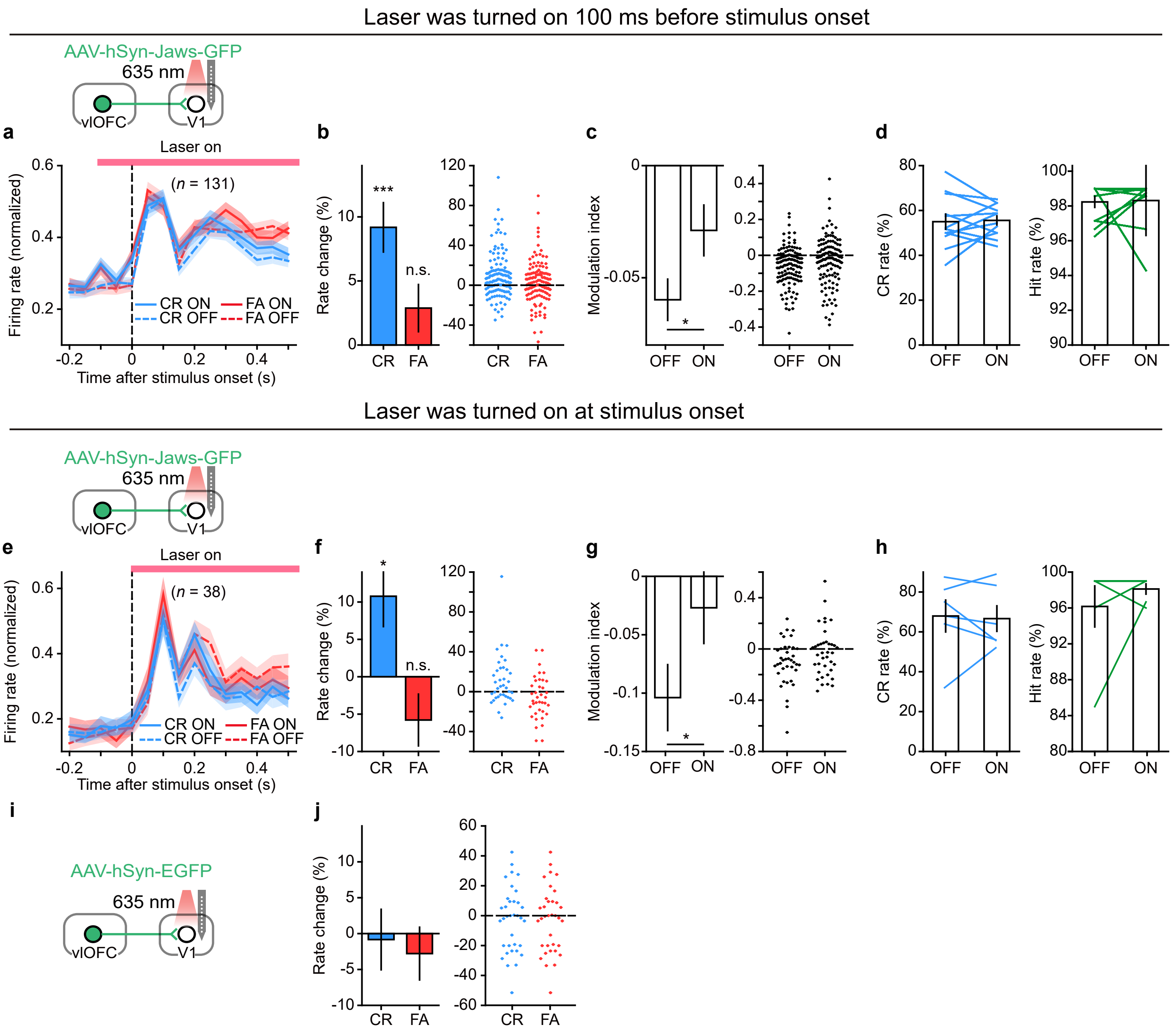


**Supplementary Figure 6 | Optogenetic inactivation of OFC projection to V1 during No-Go trials increases V1 responses to No-Go stimulus in CR condition.**

**a-d**, AAV-hSyn-Jaws-GFP was injected in the OFC. Laser stimulation in V1 during No-Go trials was turned on 100-ms before stimulus onset and covered the duration of stimulus presentation. **a**, Normalized responses to the No-Go stimulus in FA and CR trials, respectively, with and without inactivating OFC axons in V1. Horizontal line, duration of laser stimulation. **b**, Inactivating OFC axons in V1 during No-Go trials increased the responses of V1 neurons in CR (****P* = $1.7\times10^{-5}$) but not in FA trials (*P* = 0.12). *n* = 131 neurons. **c**, Inactivating OFC axons in V1 during No-Go trials significantly reduced the amplitude of MI (**P* = 0.01, *n* = 131 neurons). **d**, Inactivating OFC axons in V1 during No-Go trials did not affect the CR rate (*P* = 0.91) or hit rate (*P* = 0.77) of the mice (*n* = 12 sessions from 8 mice). **e-h**, AAV-hSyn-Jaws-GFP was injected in the OFC. Laser stimulation in V1 during No-Go trials was turned on at stimulus onset and covered the duration of stimulus presentation. **e**, Normalized responses to the No-Go stimulus in FA and CR trials, respectively, with and without inactivating OFC axons in V1. Horizontal line, duration of laser stimulation. **f**, Inactivating OFC axons in V1 during No-Go trials increased the responses of V1 neurons in CR (**P* = 0.02) but not in FA trials (*P* = 0.09). *n* = 38 neurons. **g**, Inactivating OFC axons in V1 during No-Go trials significantly reduced the amplitude of MI (**P* = 0.01, *n* = 38 neurons). **h**, Inactivating OFC axons in V1 during No-Go trials did not affect the CR rate (*P* = 0.84) or hit rate (*P* = 0.75) of the mice (*n* = 6 sessions from 6 mice). **i**, For control mice, AAV-hSyn-EGFP was injected in the OFC. Laser stimulation in V1 during No-Go trials was turned on at stimulus onset and covered the duration of stimulus presentation. **j**, For control mice, laser stimulation in V1 during No-Go trials did not cause significant change in the responses of V1 neurons in either CR (*P* = 0.72) or FA trials (*P* = 0.53). *n* = 35 neurons. Data were analyzed using Wilcoxon two-sided signed rank test. Shadings and error bars denote s.e.m. Source data are provided as a Source Data file.

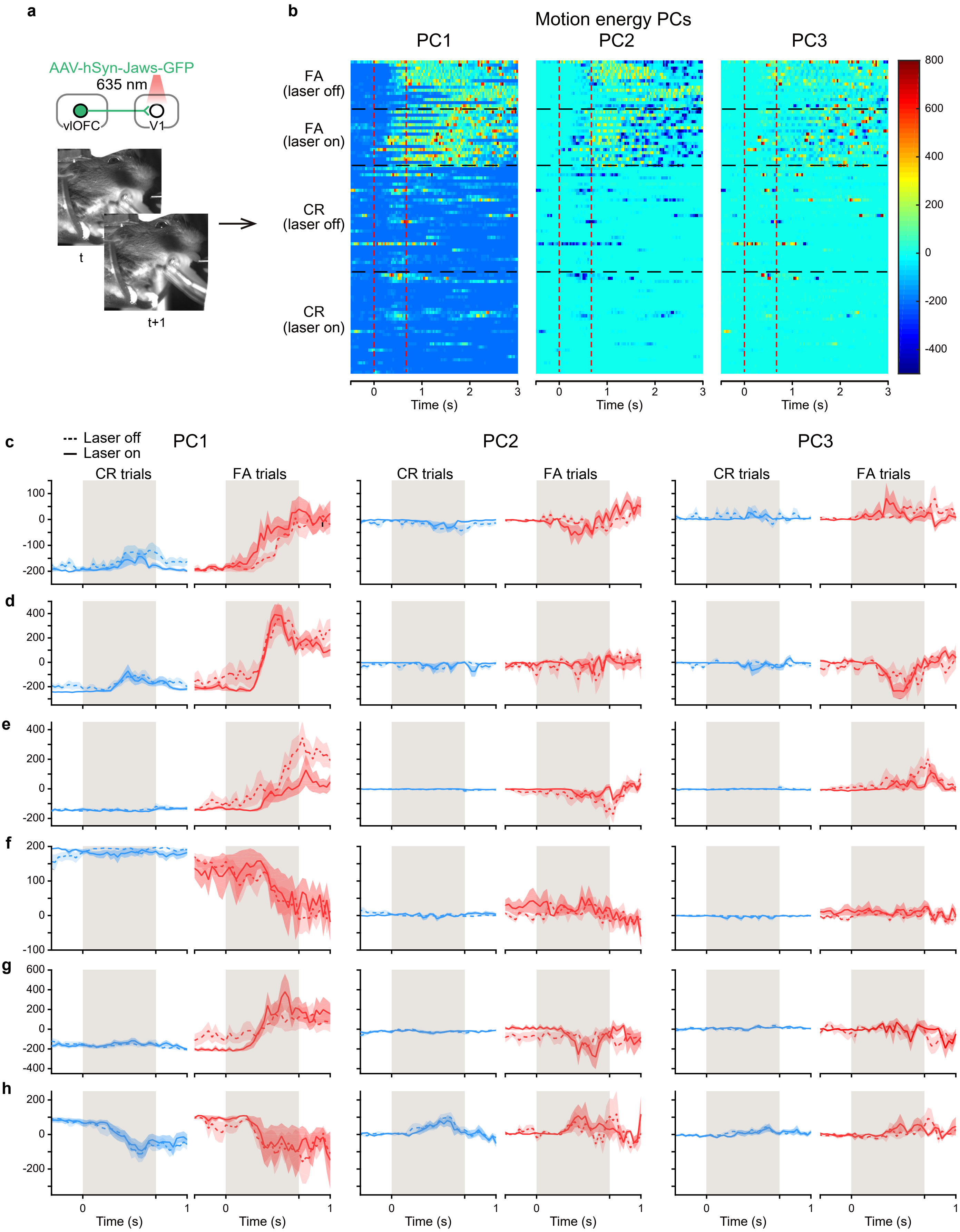
a
b
AAV-hSyn-Jaws-GFP
635 nm
vlOFC
V1
t
t+1
Motion energy PCs
PC1
PC2
PC3
FA (laser off)
FA (laser on)
CR (laser off)
CR (laser on)
Time (s)
c
d
e
f
g
h
Laser off
Laser on
CR trials
FA trials

**Supplementary Figure 7 | Optogenetic inactivation of OFC axons in V1 during No-Go trial did not affect orofacial movement during waiting period in CR trials.**

**a**, Upper, strategy for inactivating OFC projection to V1 and schematic of laser stimulation. Lower, two consecutive frames from a video recording of a mouse's face during Go/No-Go visual task. **b**, The top three principle components (PCs) of the motion energy of orofacial movement in FA (CR) trials, with or without laser stimulation during No-Go trials, from an example mouse. For each PC, the time points within the two vertical dashed lines represent the waiting period. **c**, Motion energy PC averaged across trials for the mouse shown in **b**. The gray region represents the waiting period. **d-h**, Motion energy PC averaged across trials for another 5 mice. For each of these mice, motion energy PCs in CR trials during the waiting period were not significantly different between laser off and laser on conditions. $P > 0.05$, two-way repeated measures ANOVA. The mice in **c-h** were the same as those shown in **Supplementary Fig. 6h.** The orofacial movements were analyzed using the FaceMap software (www.github.com/MouseLand/FaceMap). Shadings denote s.e.m. Source data are provided as a Source Data file.

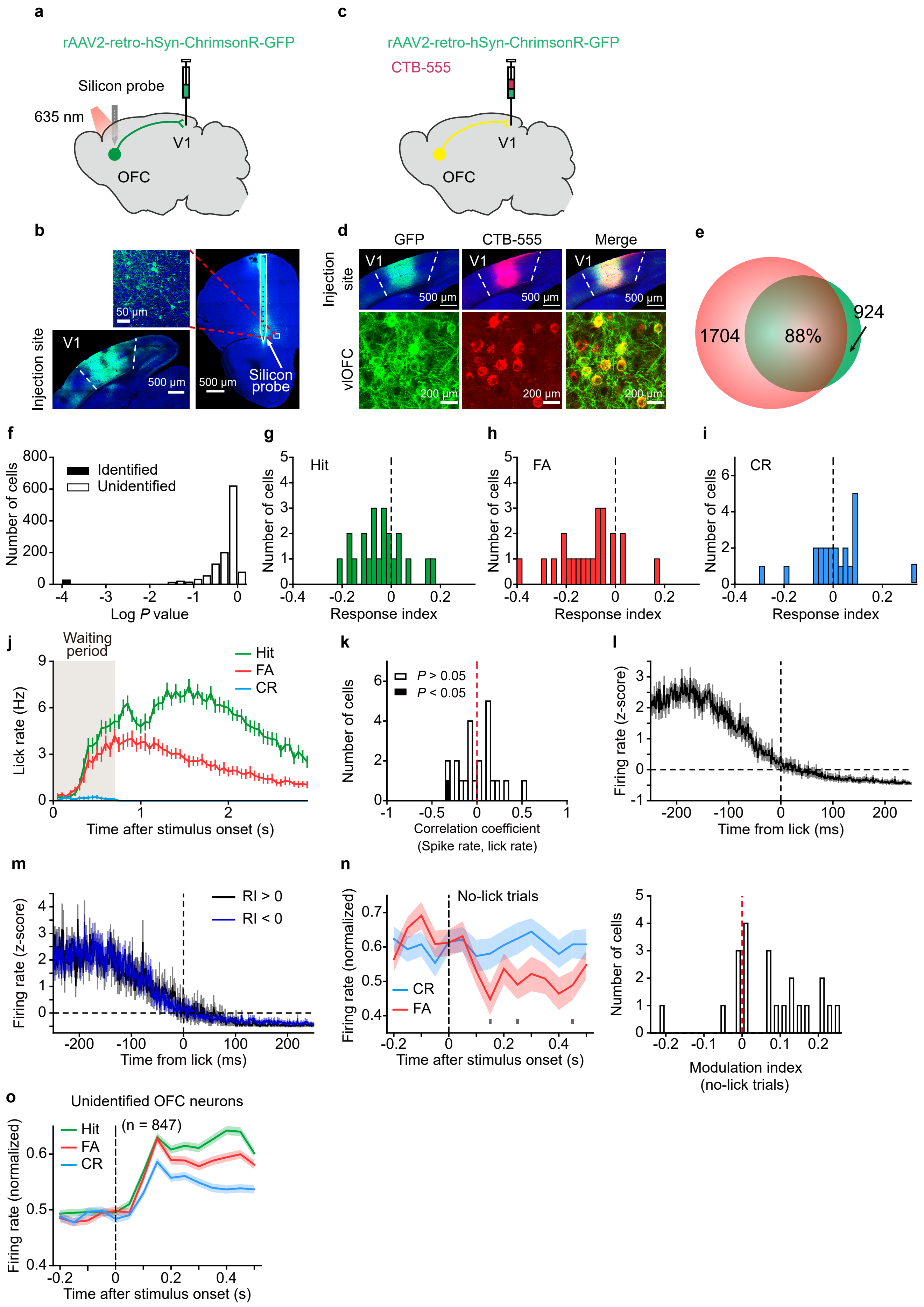
a
rAAV2-retro-hSyn-ChrimsonR-GFP
Silicon probe
635 nm
V1
OFC
c
rAAV2-retro-hSyn-ChrimsonR-GFP
CTB-555
V1
OFC
b
Injection site
V1
50 μm
500 μm
500 μm
Silicon probe
d
GFP
CTB-555
Merge
Injection site
vlOFC
V1
500 μm
200 μm
e
1704
88%
924
f
Identified
Unidentified
Number of cells
Log P value
g
Hit
Number of cells
Response index
h
FA
Number of cells
Response index
i
CR
Number of cells
Response index
j
Waiting period
Hit
FA
CR
Lick rate (Hz)
Time after stimulus onset (s)
k
P > 0.05
P < 0.05
Number of cells
Correlation coefficient
(Spike rate, lick rate)
l
Firing rate (z-score)
Time from lick (ms)
m
RI > 0
RI < 0
Firing rate (z-score)
Time from lick (ms)
n
No-lick trials
CR
FA
Firing rate (normalized)
Time after stimulus onset (s)
Number of cells
Modulation index
(no-lick trials)
o
Unidentified OFC neurons
(n = 847)
Hit
FA
CR
Firing rate (normalized)
Time after stimulus onset (s)

**Supplementary Figure 8 | Responses of identified V1-projecting OFC neurons and unidentified OFC neurons during Go/No-Go visual task.**

**a**, Schematic of rAAV2-retro-hSyn-ChrimsonR-GFP injection in V1 and phototagging in the OFC. **b**, Fluorescence images showing the injection site location in V1 (bottom-left), labeling of neurons in the ventrolateral OFC (vlOFC) and the electrode track marked by DiO (right). Enlarged view of region in white box is shown on the top-left. **c**, Schematic of co-injection of rAAV2-retro-hSyn-ChrimsonR-GFP and CTB-555 in V1. **d**, Fluorescence images showing the injection site location in V1 (upper) and labeling of neurons in the OFC (lower). Yellow shows the overlap of GFP and CTB signals in V1 (upper) or the OFC (lower). **e**, Percentage of OFC neurons co-labeled by GFP and CTB for the experiment shown in **c** and **d**, computed from the data of 3 mice. **f**, Histogram of log *P* values (derived from the SALT test) for 1175 OFC neurons recorded in the phototagging experiments. **g**-**i**, Distributions of response indexes (RIs) for identified V1-projecting OFC neurons ($n$ = 22) in hit (**g**), FA (**h**) and CR (**i**) trials. The RI was defined as $(R_{evoked} - R_{baseline})/(R_{evoked} + R_{baseline})$, where $R_{evoked}$ and $R_{baseline}$ represented the waiting-period firing rate and the baseline firing rate, respectively. The RIs in hit and FA trials were both significantly smaller than zero ($P = 0.03$ and $6.9\times10^{-4}$, respectively, Wilcoxon two-sided signed rank test). The RI in CR trials was not significantly different from zero ($P$ = 0.49, Wilcoxon two-sided signed rank test). **j**, PSTH (50 ms/bin) of licking behavior in the waiting period and the answer period for hit, FA, and CR trials, respectively ($n$ = 16 mice). The gray region indicates the waiting period. **k**, Distribution of Pearson's correlation coefficients between the spike rate of V1-projecting OFC neurons and lick rate in FA trials during the waiting period. The distribution of correlation was not significantly different from zero ($P = 0.96$, $n$ = 22, Wilcoxon two-sided signed rank test). Open and black bars denote cells with non-significant and significant coefficients, respectively. **l**, Histogram of lick-triggered spikes, computed using spikes and licks during the waiting period of all trials for V1-projecting OFC neurons. **m**, Histogram of lick-triggered spikes computed using spikes and licks within the waiting period of hit trials for V1-projecting OFC neurons (blue, RI < 0, $n$ = 6; black, RI > 0, $n$ = 13). **n**, Left, normalized responses of V1-projecting OFC neurons to the No-Go stimulus in FA and CR trials where no lick occurred during the waiting period. Gray dots, time points in which the responses between FA and CR trials were significantly different ($P < 0.05$, two-way repeated measures ANOVA $F_{(1, 21)} = 4.9$, $P = 0.039$ followed by Sidak's multiple comparisons test). Right, distribution of modulation index for V1-projecting OFC neurons, computed using those trials where no lick occurred during the waiting period ($P = 0.003$, $n$ = 22, Wilcoxon two-sided signed rank test). **o**, The responses of unidentified OFC neurons in the Go/No-Go task. The responses of each neuron were normalized by peak response across hit, FA and CR conditions, and were averaged across all neurons ($n$ = 847). The responses in FA trials were significantly higher than those in CR trials. $F_{(1, 846)} = 43.35$, $P = 8.0\times10^{-11}$, two-way repeated measures ANOVA. For **f**-**o**, source data are provided as a Source Data file. Shadings and error bars, mean ± s.e.m.

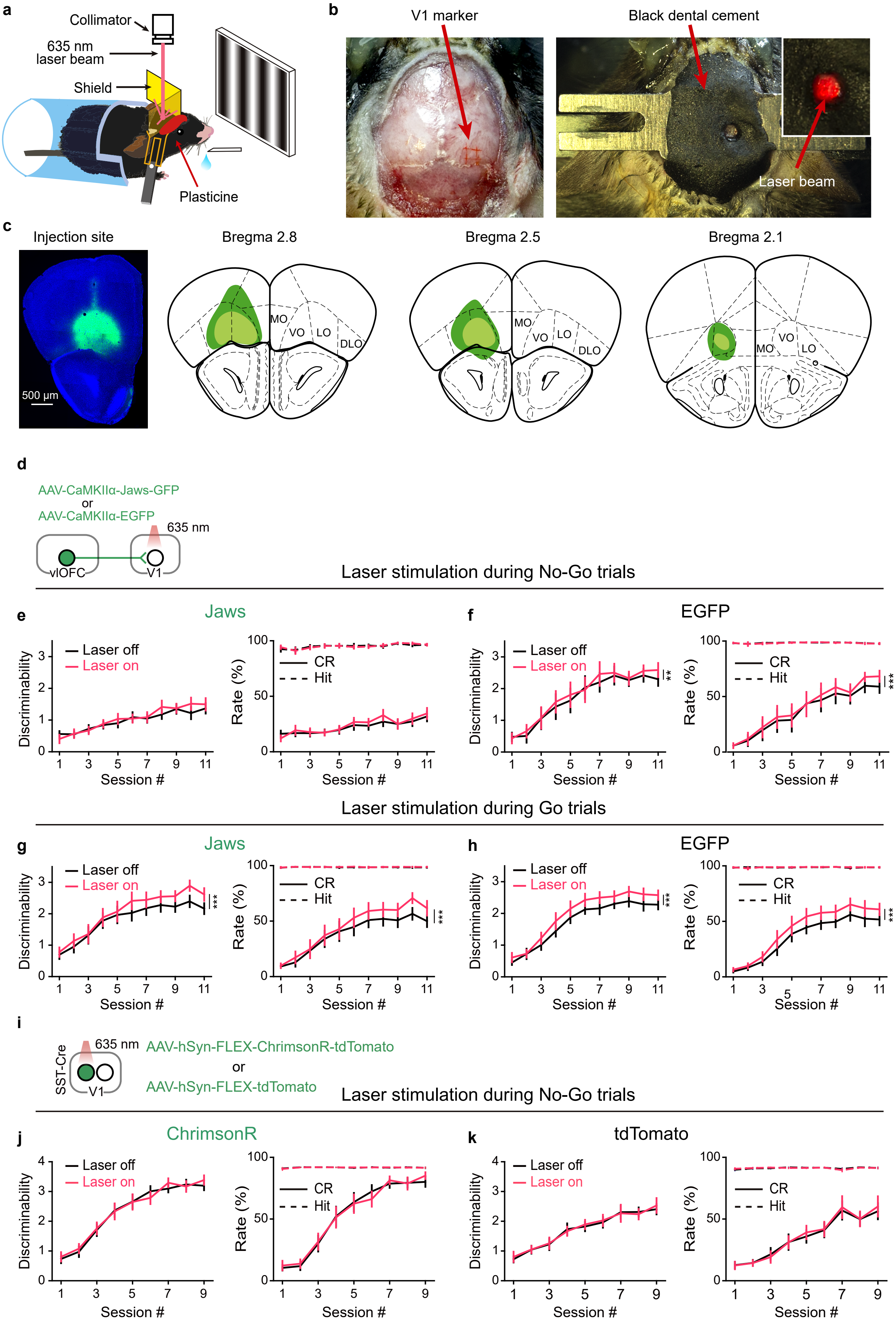

a
Collimator
635 nm laser beam
Shield
Plasticine
b
V1 marker
Black dental cement
Laser beam
c
Injection site
500 μm
Bregma 2.8
Bregma 2.5
Bregma 2.1
MO
VO
LO
DLO
d
AAV-CaMKIIα-Jaws-GFP
or
AAV-CaMKIIα-EGFP
635 nm
vlOFC
V1
Laser stimulation during No-Go trials
e
Jaws
f
EGFP
Laser off
Laser on
CR
Hit
Discriminability
Rate (%)
Session #
Laser stimulation during Go trials
g
Jaws
h
EGFP
i
SST-Cre
635 nm
V1
AAV-hSyn-FLEX-ChrimsonR-tdTomato
or
AAV-hSyn-FLEX-tdTomato
Laser stimulation during No-Go trials
j
ChrimsonR
k
tdTomato

**Supplementary Figure 9 | Effect of inactivating OFC projection to V1 or activating SST interneurons in V1 on the performance of Go/No-Go visual task.**

**a**, Images of behavioral setup and optogenetic manipulation. **b**, Left, the skull region above V1 was marked during the surgery; Right, after virus injection in the OFC the headplate was fixed to the skull using dental cement mixed with carbon powder. The cement covered the skull except the region above V1 so that photostimulation of area beyond V1 was prevented. Inset, V1 region was stimulated with red laser. **c**, Left, fluorescence image showing the expression of AAV-CaMKIIα-Jaws-GFP in the OFC in a C57BL/6 mouse. Right, the maximal (green) and minimal (light green) extent of virus expression in the OFC for all Jaws-expressing mice in **Fig. 6**. **d**, Schematic of the strategy for inactivating OFC projection to V1. **e** and **f**, Laser stimulation was applied during No-Go trials. **e**, Discriminability (left), hit and CR rates (right) with and without laser stimulation during No-Go trials for mice in which AAV-CaMKIIα-Jaws-GFP was injected in the OFC (*n* = 13). Discriminability: $F_{(1, 12)} = 1.77$, $P = 0.21$; hit rate: $F_{(1, 12)} = 0.06$, $P = 0.81$; CR rate: $F_{(1, 12)} = 3.44$, $P = 0.09$. Two-way repeated measures ANOVA. **f**, Discriminability, hit and CR rates with and without laser stimulation during No-Go trials for control mice in which AAV-CaMKIIα-EGFP was injected in the OFC (*n* = 9). Discriminability: $F_{(1, 8)} = 20.98$, $P = 0.002$; hit rate: $F_{(1, 8)} = 0.55$, $P = 0.48$; CR rate: $F_{(1, 8)} = 31.32$, $P = 5.13\times10^{-4}$. Two-way repeated measures ANOVA. **g** and **h**, Laser stimulation was applied during Go trials. **g**, Similar to those described in **e** except that laser stimulation was applied during Go trials. *n* = 12 mice in which AAV-CaMKII α-Jaws-GFP was injected in the OFC. Discriminability: $F_{(1, 11)} = 48.58$, $P = 2.36\times10^{-5}$; hit rate: $F_{(1, 11)} = 2.13$, $P = 0.17$; CR rate: $F_{(1, 11)} = 47.96$, $P = 2.5\times10^{-5}$. Two-way repeated measures ANOVA. **h**, Similar to those described in **f** except that laser stimulation was applied during Go trials. *n* = 11 mice in which AAV-CaMKIIα-EGFP was injected in the OFC. Discriminability: $F_{(1, 10)} = 38.26$, $P = 1.0\times10^{-4}$; hit rate: $F_{(1, 10)} = 1.6$, $P = 0.23$; CR rate: $F_{(1, 10)} = 44.45$, $P = 5.59\times10^{-5}$. Two-way repeated measures ANOVA. Note that for both control group (**h**) and experimental group (**g**) of mice, laser stimulation during Go trials slightly but significantly increased the discriminability and CR rate of the mice. This may be due to the possibility that laser stimulation served as a cue to guide the mice's behavior. **i**, Schematic of the strategy for activating SST interneurons in V1. **j** and **k**, Laser stimulation was applied during No-Go trials. **j**, Discriminability, hit and CR rates with and without laser stimulation during No-Go trials for SST-Cre mice in which AAV-hSyn-FLEX-ChrimsonR-tdTomato was injected in V1 (*n* = 11). Discriminability: $F_{(1, 10)} = 0.64$, $P = 0.44$; hit rate: $F_{(1,10)} = 0.05$, $P = 0.82$; CR rate: $F_{(1, 10)} = 0.32$, $P = 0.59$. Two-way repeated measures ANOVA. **k**, Similar to those described in **j** except that AAV-hSyn-FLEX-tdTomato was injected in V1 of SST-Cre mice (*n* = 10). Discriminability: $F_{(1, 9)} = 0.33$, $P = 0.58$; hit rate: $F_{(1, 9)} = 0.08$, $P = 0.78$; CR rate: $F_{(1, 9)} = 0.76$, $P = 0.41$. Two-way repeated measures ANOVA. For **e**-**h**, **j** and **k**, source data are provided as a Source Data file. Error bars, mean ± s.e.m.

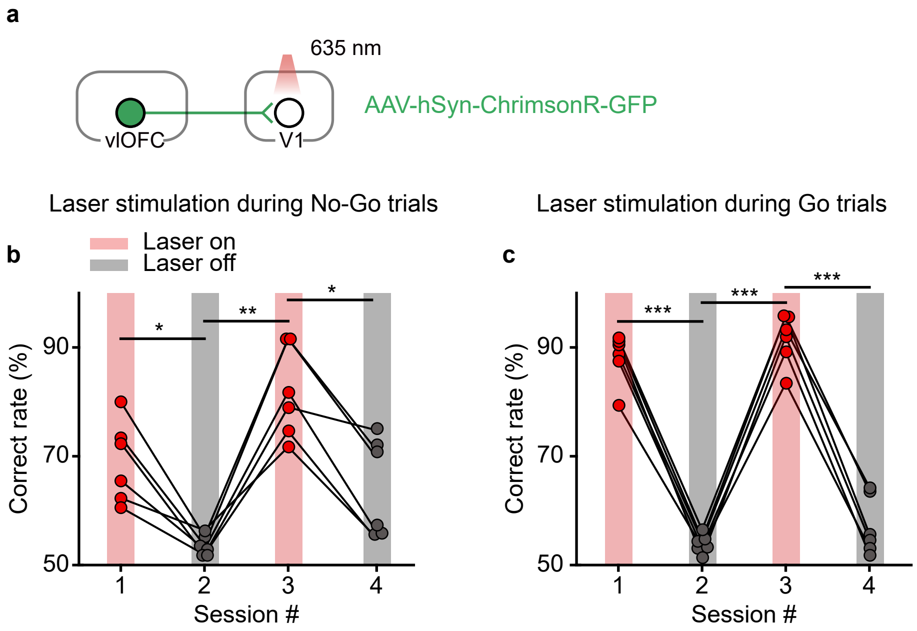


**Supplementary Figure 10 | Effect of activating OFC projection to V1 on the performance of Go/No-Go visual task.** **a**, Schematic of viral strategy to achieve optogenetic activation of OFC axons in V1. **b**, Behavioral performance of 6 mice (C57BL/6) in which laser stimulation of V1 was applied during No-Go trials in the first and third sessions. *$P < 0.05$, **$P < 0.01$, one-way repeated measures ANOVA with the Greenhouse-Geisser correction ($F_{(1.91, 9.56)} = 32.24$, $P = 6.1\times10^{-5}$) followed by Sidak's multiple comparisons test. **c**, Behavioral performance of another group of 6 mice (C57BL/6) in which laser stimulation of V1 was applied during Go trials in the first and third sessions. . ***$P < 0.001$, one-way repeated measures ANOVA with the Greenhouse-Geisser correction ($F_{(1.99, 9.94)} = 255.47$, $P = 2.9\times10^{-9}$) followed by Sidak's multiple comparisons test. Source data are provided as a Source Data file.

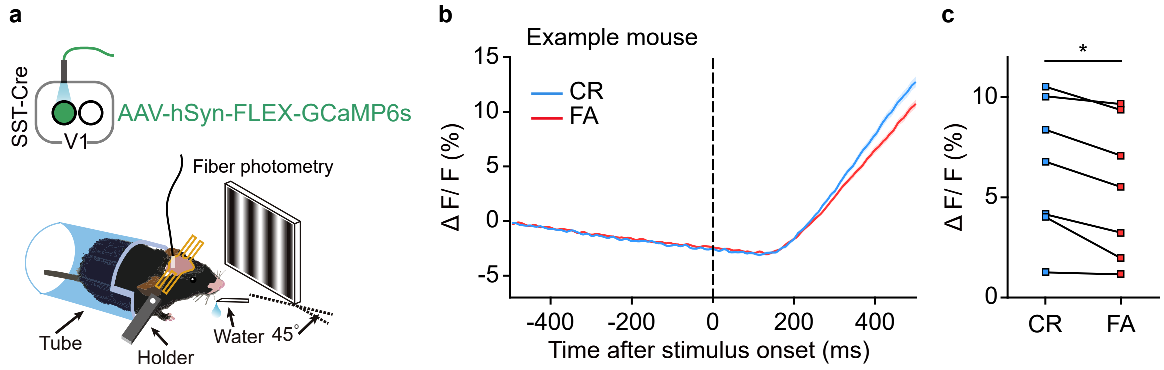


**Supplementary Figure 11 | Responses of SST interneurons in V1 to the No-Go stimulus in CR and FA trials.** **a**, Schematic of the strategy to record activities of SST interneurons in V1 from behaving mice. **b**, ΔF/F signals in FA and CR trials from an example SST-Cre mouse. Shadings, mean ± s.e.m. **c**, The responses of SST interneurons to the No-Go stimulus were significantly larger in CR than in FA trials. $P = 0.016$, $n = 7$ mice, Wilcoxon two-sided signed rank test. Source data are provided as a Source Data file.

**Supplementary Video 1 | Example of orofacial movements from trials with and without optogenetic inactivation of OFC projection to V1.**
The mouse was performing a Go/No-Go visual task. The session consisted of interleaved blocks of laser-on and laser-off trials. In laser-on blocks, laser stimulation was applied to V1 during No-Go trials to inactivate the OFC projection to V1. The video shows a hit trial, two FA trials (laser-off and laser-on), and two CR trials (laser-off and laser-on). ITI, inter-trial interval.
